# Re-evaluation of the phylogenetic diversity and global distribution of the genus *Candidatus* Accumulibacter

**DOI:** 10.1101/2021.12.20.473458

**Authors:** Francesca Petriglieri, Caitlin Singleton, Zivile Kondrotaite, Morten K. D. Dueholm, Elizabeth A. McDaniel, Katherine D. McMahon, Per H. Nielsen

## Abstract

*Candidatus* Accumulibacter was the first microorganism identified as a polyphosphate-accumulating organism (PAO), important for phosphorus removal from wastewater. This genus is diverse, and the current phylogeny and taxonomic framework appears complicated, with the majority of publicly available genomes classified as “*Candidatus* Accumulibacter phosphatis”, despite notable phylogenetic divergence. The *ppk1* marker gene allows for a finer scale differentiation into different “types” and “clades”, nevertheless taxonomic assignments remain confusing and inconsistent across studies. Therefore, a comprehensive re-evaluation is needed to establish a common understanding of this genus, both in terms of naming and basic conserved physiological traits. Here, we provide this re-assessment using a comparison of genome, *ppk1*, and 16S rRNA gene-based approaches from comprehensive datasets. We identified 15 novel species, along with the well-known *Ca*. A. phosphatis, *Ca*. A. deltensis and *Ca*. A. aalborgensis. To compare the species *in situ*, we designed new species-specific FISH probes and revealed their morphology and arrangement in activated sludge. Based on the MiDAS global survey, *Ca*. Accumulibacter species were widespread in WWTPs with phosphorus removal, indicating the process design as a major driver for their abundance. Genome mining for PAO related pathways and FISH-Raman microspectroscopy confirmed the potential for the PAO metabolism in all *Ca*. Accumulibacter species, with detection *in situ* of the typical PAO storage polymers. Genome annotation further revealed fine-scale differences in the nitrate/nitrite reduction pathways. This provides insights into the niche differentiation of these lineages, potentially explaining their coexistence in the same ecosystem while contributing to overall phosphorus and nitrogen removal.

**Importance:** *Candidatus* Accumulibacter is the most studied PAO organism, with a primary role in biological nutrient removal. However, the species-level taxonomy of this lineage is convoluted due to the use of different phylogenetic markers or genome sequencing. Here, we redefined the phylogeny of these organisms, proposing a comprehensive approach which could be used to address the classification of other diverse and uncultivated lineages. Using genome-resolved phylogeny, compared to 16S rRNA gene- and other phylogenetic markers phylogeny, we obtained a higher resolution taxonomy and established a common understanding of this genus. Furthermore, genome mining of gene and pathways of interest, validated *in situ* by application of a new set of FISH probes and Raman micromicrospectroscopy, provided additional high-resolution metabolic insights into these organisms.

## Introduction

Phosphorus (P) removal from wastewater is an essential step in wastewater treatment to prevent environmental damages (e.g., eutrophication) of receiving water bodies. The enhanced biological phosphorus removal (EBPR) process is an environmentally-friendly and cost-effective technology increasingly employed for this purpose in wastewater treatment plants (WWTPs) (1, 2). EBPR is mediated by specialized microorganisms, known as polyphosphate-accumulating organisms (PAOs), able to accumulate P as intracellular polyphosphate (poly-P), thereby allowing the removal of excess P from the water by wasting of surplus biomass (1). One of the first PAOs identified and still considered the model PAO organism, was *Candidatus* Accumulibacter from the *Rhodocyclaceae* family in the Proteobacteria (3, 4).

*Ca*. Accumulibacter species have not been isolated in pure culture, despite enrichment efforts (5, 6). Cultivation-independent approaches have been essential and widely applied to investigate these microorganisms (4, 7–12), and *Ca*. Accumulibacter populations have shown to be abundant both in lab-scale EBPR reactors (4, 13) and in full-scale EBPR plants (8, 14, 15). However, the 16S rRNA marker gene, which is a common target for culture-independent techniques is highly conserved within the genus, which prohibits the differentiation of functional important sub-genus taxa (i.e., species or strains) (16). To overcome this problem, the phylogeny of *Ca*. Accumulibacter has been resolved by sequencing of the polyphosphate kinase gene (*ppk1*) (8, 17, 18), which encodes an enzyme involved in poly-P accumulation (16). By using the *ppk1* gene, the genus has been grouped in two major divisions, called type I and type II, each with multiple subdivisions referred to as clades (clade IA-IH, IIA-III) (7, 16, 17). It has generally been assumed that this dichotomy could mirror the phenotypic variants observed under different environmental conditions, for example nitrate reduction capabilities, but this is not universally true (5, 13, 19, 20).

These discrepancies have motivated efforts to use comparative genomics to define key traits at finer scales of resolution from the retrieval of several metagenome-assembled genomes (MAGs). This was first achieved by García Martín et al., (11) with the subsequent completion of the genome for *Ca*. Accumulibacter Clade IIA strain UW-1. Recently, the application of high-throughput sequencing techniques allowed the recovery of thousands of high-quality MAGs from WWTPs ecosystems (21, 22), providing an excellent opportunity to investigate the diversity and ecophysiology of microorganisms important in these systems, including *Ca*. Accumulibacter. Genome-based approaches are also an invaluable instrument to resolve the phylogeny of this group of microorganisms. Even though *ppk1*-based phylogeny and genome-based taxonomy generally coincide, a recent study from McDaniel et al. (12) observed some discrepancies, with a few MAGs classified as *Ca*. Accumulibacter branching outside of the established taxonomy. Moreover, most of the publicly available Accumulibacter-associated genomes are currently classified as “*Candidatus* Accumulibacter phosphatis”, despite notable phylogenetic divergence, increasing the confusion in the taxonomic assignments and highlighting the need for a substantial re-evaluation of the phylogeny of this genus.

This confusion is also evident when using fluorescence *in situ* hybridization (FISH) to study the morphological diversity within this genus. Three FISH probes (PAOmix probes) were designed to target their 16S rRNA (4). However, the PAOmix has been shown to be inadequate to specifically distinguish *Ca*. Accumulibacter from other phylogenetically related taxa and it is targeting species belonging to the genus *Propionivibrio*, a well-known glycogen-accumulating organism (GAO) (23). Another FISH probe set was designed more recently to distinguish type I from type II (19) and displays a morphologically heterogeneous community. Therefore, a re-evaluation of existing FISH probes targeting *Ca*. Accumulibacter is needed for confident application in future studies. This could benefit from the use of comprehensive and ecosystem-specific full-length 16S rRNA gene reference databases, such as MiDAS3 (24, 25) and MiDAS4 (26), which facilitate the analysis of the microbial diversity in WWTPs with species-level resolution. Such databases can also be used to design a new set of species-specific FISH probes, which may subsequently be applied in combination with other techniques (e.g., Raman microspectroscopy) to confirm the metabolic potential and function of these microorganisms in the WWTP ecosystem and, finally, to supply *Candidatus* or provisional names for the key-microorganisms.

Here we used a collection of new and publicly available MAGs to obtain a comprehensive comparison of genome, *ppk1*, and 16S rRNA gene-based phylogeny to redefine the taxonomy of the *Ca*. Accumulibacter genus. Microbial community data from the MiDAS global project was used to profile the abundance and distribution of *Ca*. Accumulibacter species in full-scale WWTPs worldwide through 16S rRNA gene amplicon sequencing. The full-length 16S rRNA gene sequences were used to reevaluate existing FISH probes, and to design a set of new species-level FISH probes to determine their morphology and abundance. The FISH probes were applied in combination with Raman microspectroscopy to detect the storage polymers typical of the PAO metabolism, and these main metabolic traits were subsequently confirmed by annotation of key genes for poly-P, glycogen, and PHA accumulation as well as nitrogen metabolism. Using this approach, we identified 18 novel *Ca*. Accumulibacter species, for which we provide here *Candidatus* names, and substantially resolved the complex phylogeny of this lineage.

## Material and methods

### Sampling and fixation

Sampling of activated sludge was carried out within the Danish MiDAS project (25, 27) and the global MiDAS project (26). In short, fresh biomass samples from the aeration tank from various WWTPs were collected and either sent to Aalborg University (Danish MiDAS) or preserved in RNAlater and shipped to Aalborg University with cooling elements (Global MiDAS). Upon arrival, all samples were subsampled and stored at −20°C for sequencing workflows, and fixed for FISH with 50% ethanol (final volume) or 4% PFA (final volume), as previously described (28).

### DNA extraction and community profiling using 16S rRNA gene amplicon sequencing

DNA extraction, sample preparation and amplicon sequencing were performed as described by Nierychlo et al., (25) and Dueholm et al., (26). V1-V3 16S rRNA gene regions were amplified using the 27F (5’-AGAGTTTGATCCTGGCTCAG-3’) (29) and 534R (5’-ATTACCGCGGCTGCTGG-3’) (30) primers, and the resulting amplicons were used in all the analyses. The V4 16S rRNA gene region was amplified using the 515F (5’-GTGYCAGCMGCCGCGGTAA-3’) (31) and 806R (5’-GGACTACNVGGGTWTCTAAT-3’) (32) primers for comparison with the previous dataset. Data was analyzed using R (version 3.5.2) (33) through RStudio software (34) and visualized using ampvis2 (version 2.7.5) (35) and ggplot2 (36). Theoretical evaluation of the taxonomic resolution provided by different variable regions of the 16S rRNA gene was determined by extracting *in silico* ASV corresponding to each variable region from references in the MiDAS4 database, classifying them with the full database, and calculating the percentage of correct and wrong classifications using the MiDAS4 taxonomy as the ground truth.

### Phylogenomic analysis and MAGs annotation

MAGs identified as potential *Ca*. Accumulibacter or *Propionivibrio*, a close relative to the former, were obtained from a set of 1083 high-quality (HQ) MAGs recovered from Danish WWTPs (22). Identification was based on the GTDB-Tk v1.4.1 (37) taxonomy classification of the MAGs and mapping of extracted 16S rRNA genes against the MiDAS 3 database (22, 25) using usearch -global. MAGs with either genome taxonomy or 16S rRNA gene classification as *Ca*. Accumulibacter and *Propionivibrio* were selected for further investigation. These genomes were added to a collection of publicly available HQ MAGs (12, 21, 37–40), selected based on quality standards proposed by Bowers et al., (41) (completeness and contamination of >90% and <5%, respectively) (see **Data S1** for accessions and leaf names). The IA-UW2 strain assembled by Flowers et al. (10) was renamed to UW3, after removing a prophage contig. Concatenated and trimmed alignments of 120 single copy marker gene proteins were created using GTDB-Tk ‘classify’. The multiple sequence alignments included the selection of MAGs above as well as three *Azonexus* (former *Dechloromonas*) isolates (GCA_000012425.1, GCA_000519045.1, GCA_001551835.1) used as outgroups to create a rooted tree. The alignment was used as input for IQ-TREE v2 (42) to create a genome tree using the WAG+G model with 100x bootstrap iterations. dRep v2.3.2 (43) -comp 50 -con 10 -sa 0.95 was used to dereplicate the genomes at 95% ANI to indicate the species representatives in the genome tree. Pairwise ANI was calculated for all *Ca*. Accumulibacter genomes using fastANI (44) and ordered in the same order as the phylogenetic tree in Figure 1. MAGs and genomes were annotated as described previously (45). Briefly, EnrichM v0.5.0 (github.com/geronimp/enrichM) ‘annotate’ was used to annotate the genes with KEGG (46) Orthology (KO) numbers using DIAMOND v0.9.22 (47) BLAST against the KO annotated Uniref100 database (EnrichM database v10). EnrichM ‘classify’ with ‘--cutoff 1’ was then used to determine the presence of 100% complete KEGG modules. The output used in this study is presented in **Data File S2**. Additionally, the MAGs were uploaded to the ‘MicroScope Microbial Genome Annotation & Analysis Platform’ (MAGE) (48) in order to cross-validate KO annotations found using EnrichM.

**Figure 1.**
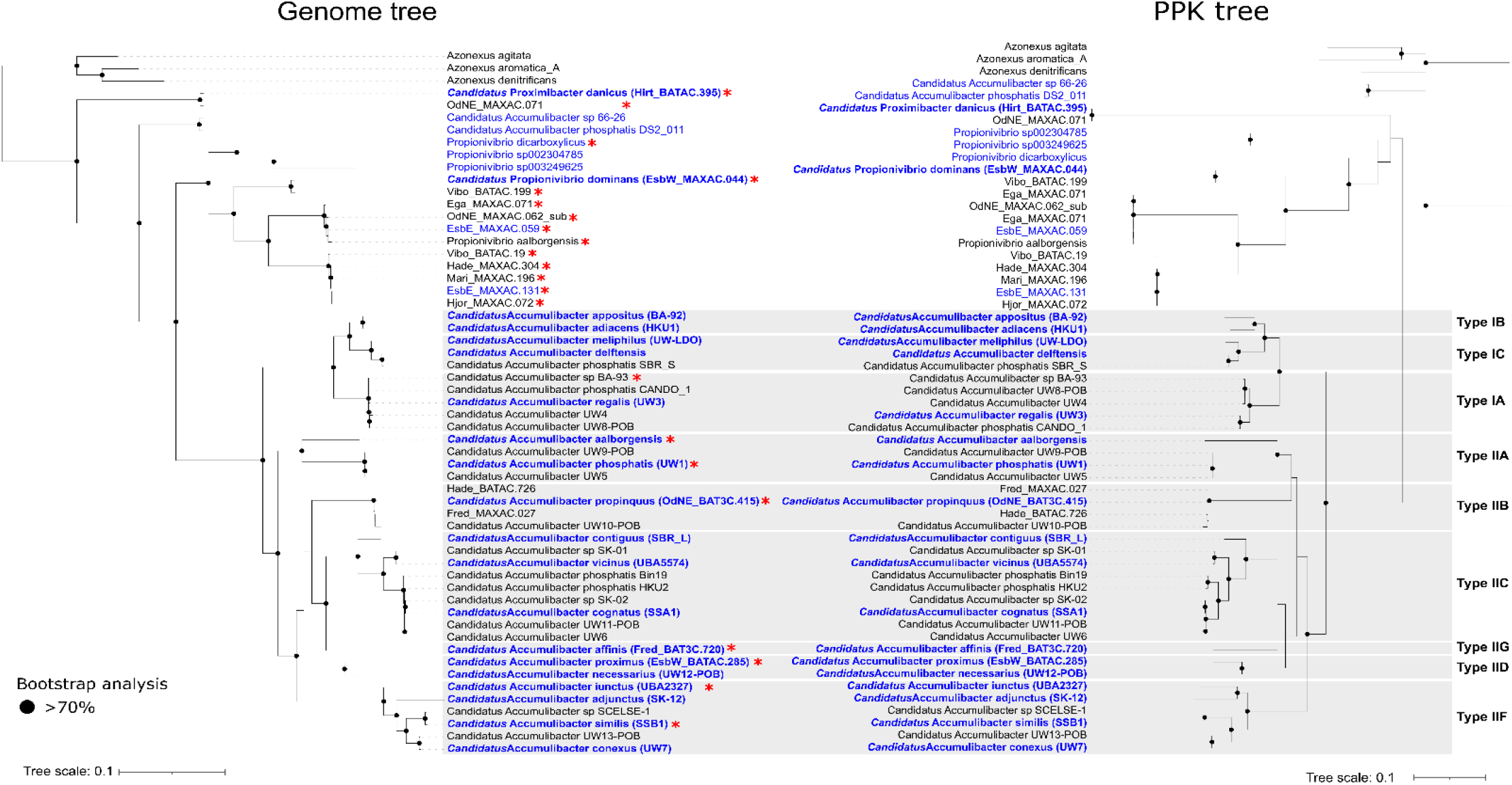
Comparison between genome- and *ppk1*-based phylogeny. The maximum likelihood genome tree was created from the concatenated alignment of 120 single copy marker gene proteins trimmed to 5000 amino acids using GTDB-Tk and 100 bootstraps. Three *Azonexus* (former *Dechloromonas*) isolates (GCA_000012425.1, GCA_000519045.1, GCA_001551835.1) were used as an outgroup. The maximum likelihood *ppk1* gene tree was created from the alignment of the *ppk1* genes extracted from the genomes, using 100 bootstraps. For NCBI GenBank genome accession numbers and leaf names see **Data S1**. Grey boxes indicate the *ppk1*-based types nomenclature. Species representatives are indicated in blue. Of these, the ones with established or proposed *Candidatus* names are indicated in bold. Red asterisk indicates MAGs with full-length 16S rRNA gene sequences.

### Ppk1 phylogenetic analysis

The *ppk1* gene sequences were sourced from the database file from McDaniel et al. (39) (github.com/elizabethmcd/ppk1_Database). Additional *ppk1* sequences from MAGs in the genome taxonomy database (GTDB) and *Propionivibrio* and *Azonexus* MAGs were sourced from the genomes (**SData 3**). Prokka v1.14 (49) was used to call and annotate the genes within the genomes, enabling identification of the polyphosphate kinase (*ppk1*), which were extracted using Fxtract v2.3 (github.com/ctSkennerton/fxtract) and added to the *ppk1* database file. The *ppk1* gene sequences were aligned using MAFFT v7.47 (50) with the mafft –auto command. The alignment was input into IQ-TREE v2 with Model Finder enabled using ‘-m MFP’. The GTR+F+I+G4 model was selected by Model Finder, and a phylogenetic maximum likelihood tree was created with 100x bootstrap iterations. ARB v6.0.3 (51) was used to visualise the trees and set the root based on the outgroup sequences. The trees were exported to iTOL v6 (52), enabling the nodes to be matched up as much as possible for presentation in **Figure 1**. Final aesthetic processing was done in Inkscape v1.0.2.

### 16S rRNA gene phylogenetic analysis, FISH probes design and evaluation

Phylogenetic analysis of 16S rRNA gene sequences and design of FISH probes were performed using the ARB software v.6.0.6 (51). 16S rRNA gene sequences were extracted from the MAG gene files created by prokka (.ffn files) using Fxtract and were also retrieved from the MiDAS4 database (26) and a publicly available set (3). A phylogenetic tree was calculated based on comparative analysis of aligned 16S rRNA gene sequences using the maximum likelihood method and a 1000 – replicates bootstrap analysis. Coverage and specificity of the FISH probes were evaluated and validated *in silico* with the MathFISH software for hybridization efficiencies of target and potentially weak non-target matches (53). When needed, unlabelled competitors were designed. All probes were purchased from Biomers (Ulm, Germany), labelled with cyanine 3 (Cy3), cyanine 5 (Cy5), FLUOS, Atto 532, Atto 565, Atto 594, and Atto 633 fluorochromes.

### FISH, quantitative FISH (qFISH), and Raman microspectroscopy

FISH was performed as previously described (28). Optimal formamide concentration for FISH probes was determined after performing hybridization at different formamide concentrations in the range 0-70% (with 5% increments). The intensity of at least 50 cells was measured using ImageJ (54) software. Optimal hybridization conditions are described in **Table S1**. EUBmix (55, 56) was used to target all bacteria and NON-EUB (57) was used as a negative control for sequence independent probe binding. Quantitative FISH (qFISH) biovolume fractions of individual genera were calculated as a percentage area of the total biovolume, hybridizing with both EUBmix probes and a specific probe. qFISH analyses, performed using the Daime image analysis software (58), were based on 30 fields of view taken at 63× magnification. Microscopic analysis was performed with Axioskop epifluorescence microscope (Carl Zeiss, Germany) equipped with LEICA DFC7000 T CCD camera or a white light laser confocal microscope (Leica TCS SP8 X). Multicolor FISH was performed as described by Lukumbuzya et al., (59). Raman microspectroscopy combined with FISH was used to detect intracellular storage polymers (poly-P, PHA and glycogen) in probe-defined species, and was performed as previously described (60).

## Results and discussion

### Re-evaluation of the phylogeny of the genus *Ca.* Accumulibacter and other related *Rhodocyclaceae* family members

Seventeen MAGs with either GTDB taxonomy or MiDAS3 16S rRNA gene classification as *Ca*. Accumulibacter or *Propionivibrio* were identified in a set of 1083 HQ MAGs recovered from Danish WWTPs (22). These genomes were added to a collection of publicly available MAGs (12, 21, 37–40) meeting the completeness and contamination quality thresholds for HQ MAGs (>90% completeness, <5% contamination) to obtain a comprehensive overview of the phylogenetic relationship between known *Ca*. Accumulibacter and related *Rhodocyclaceae* family members, and to resolve the classification of these genera. Phylogenomic analysis based on conserved marker genes (**Figure 1**) revealed a distinction into several different genera: *Ca*. Accumulibacter, *Propionivibrio, Azonexus* (formerly *Dechloromonas*), and a previously undescribed genus. A total of 36 MAGs retrieved from complex communities, often representing a mixture of several strains (61), clustered within the genus *Ca*. Accumulibacter and, based on the proposed genome-wide ANI cutoff of 95% for species (62, 63), we identified representatives for 18 species (**Figure 1**). Only two of these matched the previously described species *Ca*. A. aalborgensis and *Ca*. A. delftensis. While GTDB-based taxonomy recognized many of the MAGs as different species, it still identified the majority as “*Candidatus* Accumulibacter phosphatis”, with an appended letter to distinguish them, because of the lack of proposed names (**Figure 1, Figure S1**).

In the first instance, we proposed *Candidatus* names to the species identified based on genome-phylogeny which were also linked to full-length 16S rRNA gene sequences. Of the 18 species representative MAGs, only six possessed full-length 16S rRNA gene sequences. Among these, we propose the MAG Candidatus Accumulibacter phosphatis UW1 (11) as the genus representative, as the highest quality and first MAG retrieved for this genus, and we assigned the species name *Candidatus* Accumulibacter phosphatis. For the remaining five species, we propose the names, *Candidatus* Accumulibacter propinquus, *Candidatus* Accumulibacter affinis, *Candidatus* Accumulibacter proximus, *Candidatus* Accumulibacter iunctus and *Candidatus* Accumulibacter similis (**Figure 1, Table 1**). However, as the genome- and *ppk1*-based phylogeny were largely concordant (**Figure 1, Figure S1**), we can confidently propose names for the remaining species representatives despite the lack of the 16S rRNA gene sequence in the MAGs (**Figure 1, Table 1**).

**Table 1.** Different phylogenetic taxonomies of species in *Ca*. Accumulibacter and related genera. MAG identifier as shown in the genome tree, *ppk*-based classification, MiDAS4-classification with ASV details, and new names proposed in this study are shown. “-”: not applicable.

| MAG identifier | PPK-based classification | MiDAS4-classification and ASV details (100% identity) | Proposed species name |
| --- | --- | --- | --- |
| *Candidatus* Accumulibacter phosphatis UW3 | IA | - | *Ca. Accumulibacter regalis* UW3 |
| *Candidatus* Accumulibacter sp BA-93 | IA | - | *Ca. Accumulibacter regalis* BA-93 |
| *Candidatus* Accumulibacter phosphatis CANDO_1 | IA | - | *Ca. Accumulibacter regalis* CANDO_1 |
| *Candidatus* Accumulibacter UW4 | IA | - | *Ca. Accumulibacter regalis* UW4 |
| *Candidatus* Accumulibacter UW8-POB | IA | - | *Ca. Accumulibacter regalis* UW8-POB |
| *Candidatus* Accumulibacter sp BA-92 | IB | - | *Ca. Accumulibacter appositus* BA-92 |
| *Candidatus* Accumulibacter phosphatis HKU1 | IB | - | *Ca. Accumulibacter adiacens* HKU1 |
| *Candidatus* Accumulibacter UW-LDO | IC | - | *Ca. Accumulibacter meliphilus* UW-LDO |
| *Candidatus* Accumulibacter delftensis | IC | *Ca. Accumulibacter aalborgensis* (ASV402, ASV1099, ASV1223, ASV2139) | *Ca. Accumulibacter delftensis* |
| *Candidatus* Accumulibacter phosphatis SBR_S | IC | - | *Ca. Accumulibacter delftensis* SBR_S |
| *Candidatus* Accumulibacter aalborgensis | IIA | *Ca. Accumulibacter aalborgensis* (ASV771, ASV2139) | *Ca. Accumulibacter aalborgensis* |
| *Candidatus* Accumulibacter phosphatis UW1 | IIA | *Ca. Accumulibacter phosphatis* (ASV402, ASV1099, ASV1223, ASV2139) | *Ca. Accumulibacter phosphatis* UW1 |
| *Candidatus* Accumulibacter UW9-POB | IIA | - | *Ca. Accumulibacter phosphatis* UW9-POB |
| *Candidatus* Accumulibacter UW5 | IIA | - | *Ca. Accumulibacter phosphatis* UW5 |
| OdNE_BAT3C.415 | IIB | *Ca. Accumulibacter phosphatis* (ASV548, ASV865) | *Ca. Accumulibacter propinquus* |
| Hade_BATAC.726 | IIB | *Ca. Accumulibacter phosphatis* (ASV548, ASV865) | *Ca. Accumulibacter propinquus* |
| Fred_MAXAC.027 | IIB | *Ca. Accumulibacter phosphatis* (ASV548, ASV865) | *Ca. Accumulibacter propinquus* |
| *Candidatus* Accumulibacter UW10-POB | IIB | - | *Ca. Accumulibacter propinquus* UW10-POB |
| *Candidatus* Accumulibacter phosphatis SBR_L | IIC | - | *Ca. Accumulibacter contiguus* SBR_L |
| *Candidatus* Accumulibacter sp SK-01 | IIC | - | *Ca. Accumulibacter vicinus* SK-01 |
| Accumulibacter phosphatis_E UBA5574 | IIC | - | *Ca. Accumulibacter vicinus* UBA5574 |
| *Candidatus* Accumulibacter phosphatis Bin19 | IIC | - | *Ca. Accumulibacter cognatus* Bin19 |
| *Candidatus* Accumulibacter phosphatis HKU2 | IIC | - | *Ca. Accumulibacter cognatus* HKU2 |
| *Candidatus* Accumulibacter sp SK-02 | IIC | - | *Ca. Accumulibacter cognatus* SK-02 |
| *Candidatus* Accumulibacter phosphatis SSA1 | IIC | - | *Ca. Accumulibacter cognatus* SSA1 |
| *Candidatus* Accumulibacter UW11-POB | IIC | - | *Ca. Accumulibacter cognatus* UW11-POB |
| *Candidatus* Accumulibacter UW6 | IIC | - | *Ca. Accumulibacter cognatus* UW6 |
| EsbW_BATAC.285 | IID | *Ca. Accumulibacter phosphatis* (ASV548) | *Ca. Accumulibacter proximus* |
| *Candidatus* Accumulibacter UW12-POB | IID | - | *Ca. Accumulibacter necessarius* UW12-POB |
| *Candidatus* Accumulibacter_B UBA2327 | IIF | *Ca. Accumulibacter midas_s_12920* (ASV908) | *Ca. Accumulibacter iunctus* UBA2327 |
| *Candidatus* Accumulibacter sp SK-12 | IIF | - | *Ca. Accumulibacter adjunctus* SK-12 |
| *Candidatus* Accumulibacter sp SCELSE-1 | IIF | - | *Ca. Accumulibacter similis* SCELSE-1 |
| *Candidatus* Accumulibacter phosphatis SSB1 | IIF | *Ca. Accumulibacter midas_s_12920* (ASV908) | *Ca. Accumulibacter similis* SSB1 |
| *Candidatus* Accumulibacter UW13-POB | IIF | - | *Ca. Accumulibacter conexus* UW13-POB |
| *Candidatus* Accumulibacter UW7 | IIF | - | *Ca. Accumulibacter conexus* UW7 |
| Fred_BAT3C.720 | IIG | *Ca. Accumulibacter phosphatis* (ASV548) | *Ca. Accumulibacter affinis* |
| Hirt_BATAC.395 | N/A | *Ca. Accumulibacter midas_s_168* (ASV154, ASV471) | *Ca. Proximibacter danicus* |
| EsbW_MAXAC.044 | N/A | *Ca. Accumulibacter midas_s_315* (ASV124, ASV600) | *Ca. Propionivibrio dominans* |

The genome-resolved phylogeny broadly mirrored the “types” division based on *ppk1* phylogeny commonly used for the *Ca*. Accumulibacter genus (**Figure 1**). According to the ANI analysis (**SFigure S1**), *Ca*. Accumulibacter MAGs within individual *ppk1*-defined clades fell within the >95% ANI cut off, while types I and II genomes were similar by approximately 80-85% ANI, as recently observed by McDaniel et al. (12). Based on these results and the proposed genus ANI boundary of 75-77% (62, 63), there is no evidence supporting the division of type I and type II genomes in separate genera. However, the dichotomy into the two types seem to indicate a phylogenetically relevant clustering into clades, and could still be useful for future studies to define different clusters at the inter-species level.

When present, the 16S rRNA gene sequences from the MAGs were mapped against the MiDAS4 full-length ASVs database, which showed that the 16S rRNA gene were not able to resolve all genome-inferred species in the genus (**Table 1, Figure 2**). The 16S rRNA gene sequences extracted from the MAGs represented members of the MiDAS-defined species *Ca*. Accumulibacter phosphatis (4 MAGs), *Ca*. Accumulibacter aalborgensis (2 MAGs) and the *de novo* species midas_s_12920 (2 MAGs). This lack of taxonomic resolution could be explained by the more rapid evolution of the *ppk1* as compared to the 16S rRNA gene, which is more conserved within the genus. The lack of resolution was also observed when analysing 16S rRNA gene sequences across the ∼500 bp amplicon sequence variants (V1-V3 region), normally used for abundance estimation (**Table 1**). Among the most abundant *Ca*. Accumulibacter ASVs in the MiDAS4 global WWTP dataset (**Figure S2**), ASV402 was 100% identical to the V1-V3 regions of both *Ca*. A. phosphatis and *Ca*. A. delftensis, ASV548 was identical to *Ca*. A. affinis, *Ca*. A. proximus, and *Ca*. A. propinquus, complicating the interpretation of amplicon abundance studies based on 16S rRNA gene amplicon sequencing.

**Figure 2.**
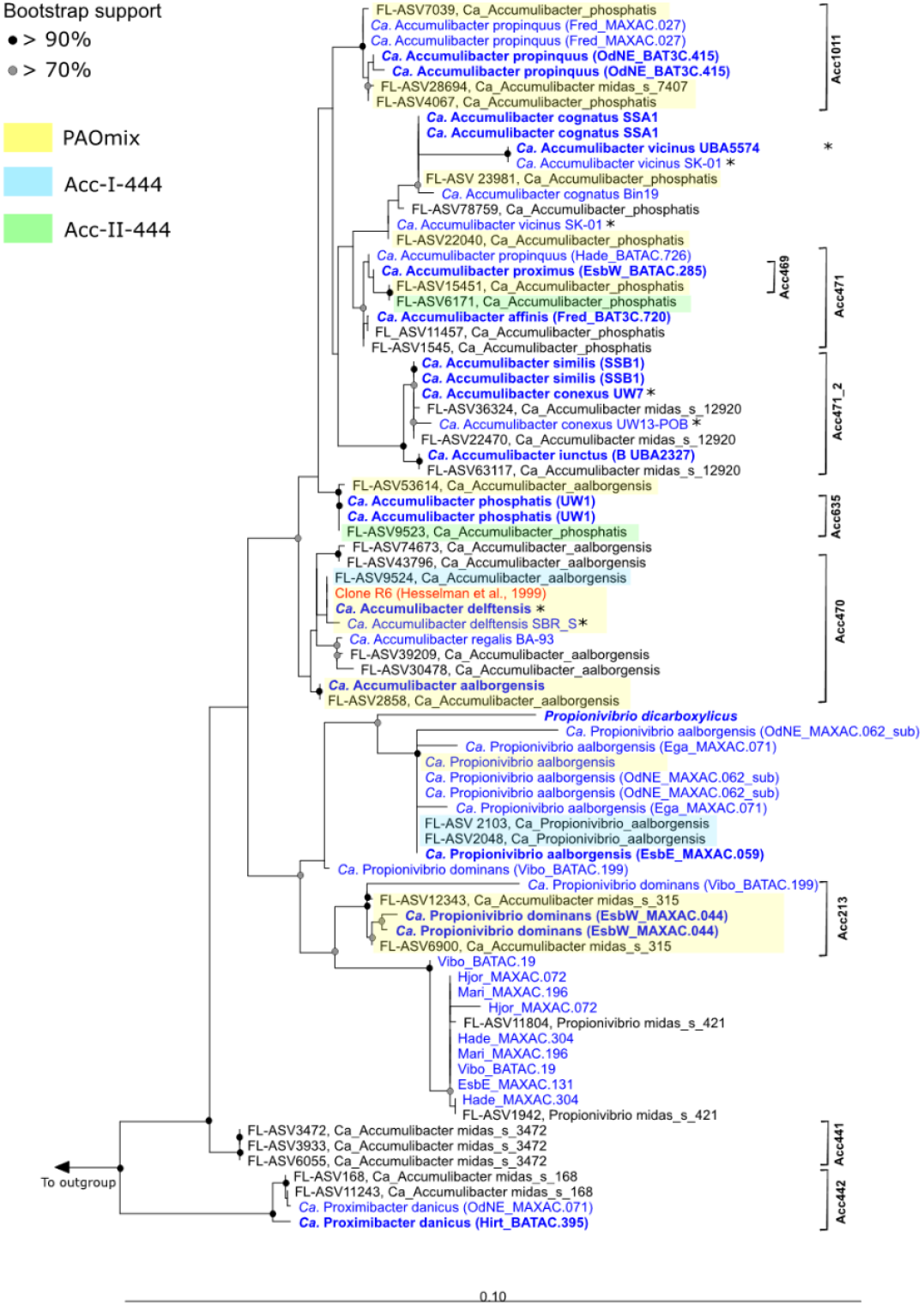
Maximum-likelihood (PhyML)16S rRNA gene phylogenetic tree of *Ca*. Accumulibacter and related species. 16S rRNA gene sequences retrieved from the MAGs are indicated in blue, the original 16S rRNA gene sequence (AJ224937) retrieved from Hesselman et. al. (3) is indicated in red. The species renamed in this study are indicated in bold blue. 16S rRNA gene partial sequences are indicated with a black asterisk. The alignment used for the tree applied a 20% conservational filter to remove hypervariable positions, giving 1250 aligned positions. Coverage of the FISH probes designed in this study is indicated with black brackets and is based on the MiDAS4 database (26). Probe coverage of widely applied probes for the *Ca*. Accumulibacter clades is shown with yellow (PAO651), orange (Acc-I-44) and red boxed (Acc-II-444). Bootstrap values from 1000 resamplings are indicated for branches with >70% (gray dot) and >90% (black) support. Species of the genus *Dechloromonas* were used as the outgroup. The scale bar represents substitutions per nucleotide base.

The MiDAS4 database presented three more *de novo* species classified as *Ca*. Accumulibacter: midas_s_315, midas_s_168, and midas_s_3472 based on full-length 16S rRNA genes. According to genome-based phylogeny (**Figure 1**), midas_s_315 clustered with the isolate genome *Propionivibrio dicarboxylicus* (GCA_900099695) and *Ca*. Propionivibrio aalborgensis MAG (GCA_900089945), together with another novel species with provisional name midas_s_421. We propose the name *Candidatus* Propionivibrio dominans for midas_s_315. The species midas_s_168 was represented by two MAGs, which clustered outside of both *Ca*. Accumulibacter and *Propionivibrio*, representing an undescribed genus and species, for which we propose the name *Candidatus* Proximibacter danicus. Due to the lack of a representative MAG, no confident taxonomic assignment could be made for the species midas_s_3472, which was investigated *in situ* (see below) to examine its capacity to carry out canonical PAO metabolism. This imprecise naming may be accounted for by the naive taxonomic assignment of the automated 16S rRNA-based taxonomy assignments with AutoTax which uses a strict species identity cut-off of 98.7% (24), which was shown to be less suited for *Ca*. Accumulibacter due to the high conservation of the 16S rRNA gene within this genus.

### Geographic distribution of *Ca.* Accumulibacter populations in global WWTPs

Despite the lack of resolution of 16S rRNA gene amplicon studies for these lineages, the new MiDAS global reference database of full-length 16S rRNA gene sequences (26) from 847 samples from 480 WWTPs located in 30 countries, ensure that amplicon analysis is still a powerful tool to analyse their geographical distribution. On a global scale, MiDAS-defined *Ca*. A. phosphatis, *Ca*. A. aalborgensis, and the *de novo* midas_s_12920, were the most abundant species within the *Ca*. Accumulibacter lineage (**Figure 3A-B**). The distribution of *Ca*. Accumulibacter was strongly influenced by the activated sludge process configuration, with higher relative abundances of all the species in EBPR plants (**Figure 3A**). MiDAS-defined *Ca*. Accumulibacter species were present in several full-scale EBPR plants worldwide, with the highest relative abundance in Mexico (5.7%), Italy (5.5%), Canada (3.9%), and South Africa (2.5%) (**Figure 3B**). Also *Ca*. Propionivibrio dominans (midas_s_315) was observed in higher abundance in BNR (biological nutrient removal) plants with nitrogen and phosphorus removal (**Figure 3A**), along with other *Propionivibrio* species (**Figure S3A**), most likely favoured by their glycogen-accumulating metabolism which can exploit the resources in the alternating aerobic/anaerobic cycling typical of the EBPR design. The highest relative abundances were observed in Norway (2.6%), the Netherlands (1.7%), and Sweden (1.2%) (**Figure S3B**). *Ca*. Proximibacter danicus was present also in WWTPs with simpler designs (**Figure S4A**). The highest abundance was observed in the United Kingdom (1.2%), Germany (0.3%), and the Czeck Republic (0.2%) (**Figure S4B**).

**Figure 3.**
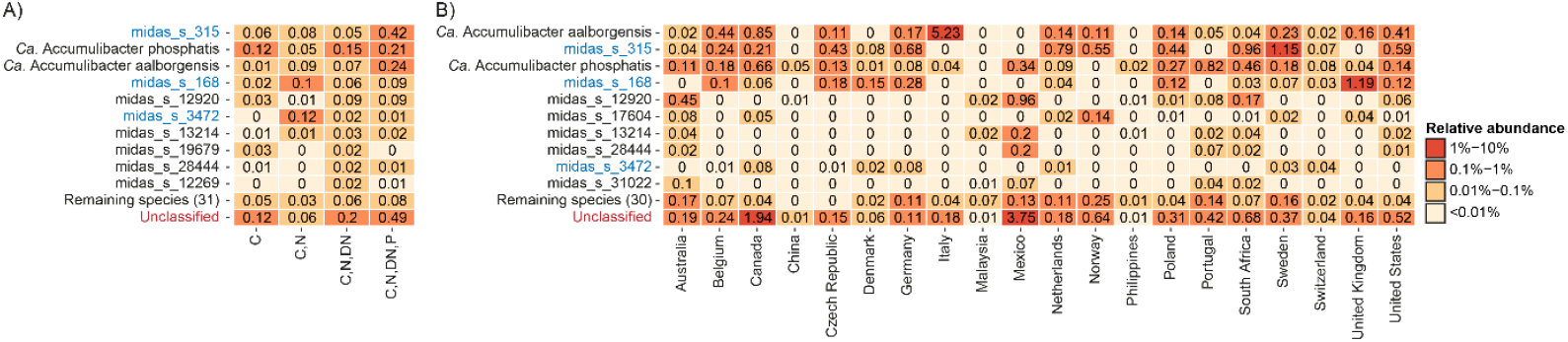
The ten most abundant *Ca*. Accumulibacter species worldwide according to the MiDAS4 survey. (A) Mean relative abundance across different process configurations (C - carbon removal; N – nitrification; DN - denitrification; P - phosphorus removal). (B) Mean relative abundance in EBPR plants across different across different countries. Data originates from the global survey of microbial communities in WWTPs (Dueholm et al., 2021) and it is based on V1-V3 amplicon dataset. Species marked in blue were wrongly classified as *Ca*. Accumulibacter according to the genome-based taxonomy.

The choice of primers has great importance for microbial profiling of activated sludge samples (64). According the MiDAS global study, *Ca*. Accumulibacter was equally well detected at the genus-level with primers targeting the V1-V3 and V4 variable regions of the 16S rRNA gene (26). This is also in accordance to what was recently observed by Roy et al., (65). However, there may be clear differences in how good amplicons from different variable regions are in resolving the species-level diversity within specific genera (24) We therefore used the MiDAS4 reference database to determine the ecosystem-specific theoretical taxonomic resolution provided by amplicons targeting different variable regions of the 16S rRNA gene (**Figure S5A)**. This revealed that amplicons targeting the V1-V3 were much better suited for resolving the species-level diversity within *Ca*. Accumulibacter compared to amplicons targeting the popular V4 regions. This was also clear when we examined the global diversity of *Ca*. Accumulibacter species based on V4 amplicon data from the MiDAS global survey, which revealed that almost all ASVs were unclassified at the species-level (**Figure S5B**). These remarkable differences must be taken into consideration when comparing abundance estimates obtained with different primer sets, but also if EBPR efficiency is evaluated using amplicon sequencing data.

### In situ visualization of *Ca.* Accumulibacter and other related species

Using the comprehensive set of ASV-resolved full-length 16S rRNA genes in the MiDAS4 database, we designed, when possible, species-specific FISH probes (**Figure 2, Table S1**) to visualize the new *Ca*. Accumulibacter and related species. When targeting *Ca*. A. phosphatis, *Ca*. A. affinis, *Ca*. A. proximus, *Ca*. A. propinquus, *Ca*. A. iunctus, and *Ca*. A. similis. *Ca*. Propionivibrio dominans the FISH probes hybridized with cocci of different diameters always arranged in small clusters inside the activated sludge floc (**Table 2, Figure 4, Figure S7**). *Ca*. Proximibacter danicus were in contrast found as rod-shaped cells often attached to filamentous bacteria as epiphytic growth (**Table 2, Figure S6**). A FISH probe designed to target the *de novo* species midas_s_3472 hybridized with low abundant, rod-shaped cells, dispersed into the floc (**Table 2, Figure S6**). The newly designed FISH probes were also applied to Danish and, when possible, to global activated sludge samples for quantitative FISH (**Table S2**). Compared to amplicon sequencing abundances, the FISH quantification provides an independent quantification based on biovolume of the specific *Ca*. Accumulibacter species. The species *Ca*. A. regalis, *Ca*. A. affinis, and *Ca*. A. proximus were abundant (>1%) in samples with high read abundance of the 16S MiDAS-defined *Ca*. Accumulibacter. The MiDAS-defined *Ca*. A. aalborgensis (which also covers *Ca*. A. delftensis) was also present in high abundance in the samples analysed. The FISH-based abundance estimates of *Ca*. A. iunctus were significantly lower than expected based on amplicon sequencing (**Table S2**), perhaps due to their small biovolume per cell. Also *Ca*. P. dominans, *Ca*. P. danicus, and the *de novo* species midas_s_3472 were generally present, but in low abundance.

**Table 2.** Summary table of the features of the different *Ca*. Accumulibacter species.

| Identifier name | FISH Probe | Morphology<br>(diameter × length in µm) | Storage polymers* |  |  |
| --- | --- | --- | --- | --- | --- |
|  |  |  | poly-P | PHA | Glycogen |
| *Ca. Accumulibacter delftensis* | Acc470 | - | + | + | + |
| *Ca. Accumulibacter regalis* | Acc635 | Coccoid (0.4-0.6) | + | + | + |
| *Ca. Accumulibacter aalborgensis* | Acc470 | Coccoid (0.7-0.9) | + | + | + |
| *Ca. Accumulibacter propinquus* | Acc1011 | Coccoid (0.8-1.2) | + | + | + |
| *Ca. Accumulibacter affinis* | Acc471 | Coccoid (0.5-0.7) | + | + | + |
| *Ca. Accumulibacter proximus* | Acc471 | Coccoid (0.5-0.7) | + | + | + |
| *Ca. Accumulibacter iunctus* | Acc471_2 | Coccoid (0.8-0.9) | + | + | + |
| *Ca. Accumulibacter similis* | Acc471_2 | Coccoid (0.8-0.9) | + | + | + |
| *Ca. Proximbacter danicus* | Acc442 | Rod-shaped (0.3-0.5 × 1-2) | - | + | - |
| *Ca. Propionivibrio dominans* | Acc213 | Rod-shaped (0.5-0.6 × 0.9-1.1) | - | + | + |
| midas_s_3472 | Acc441 | Rod-shaped (0.3-0.4 × 0.8-1.2) | - | + | + |

\*detected by Raman microspectroscopy

**Figure 4.**
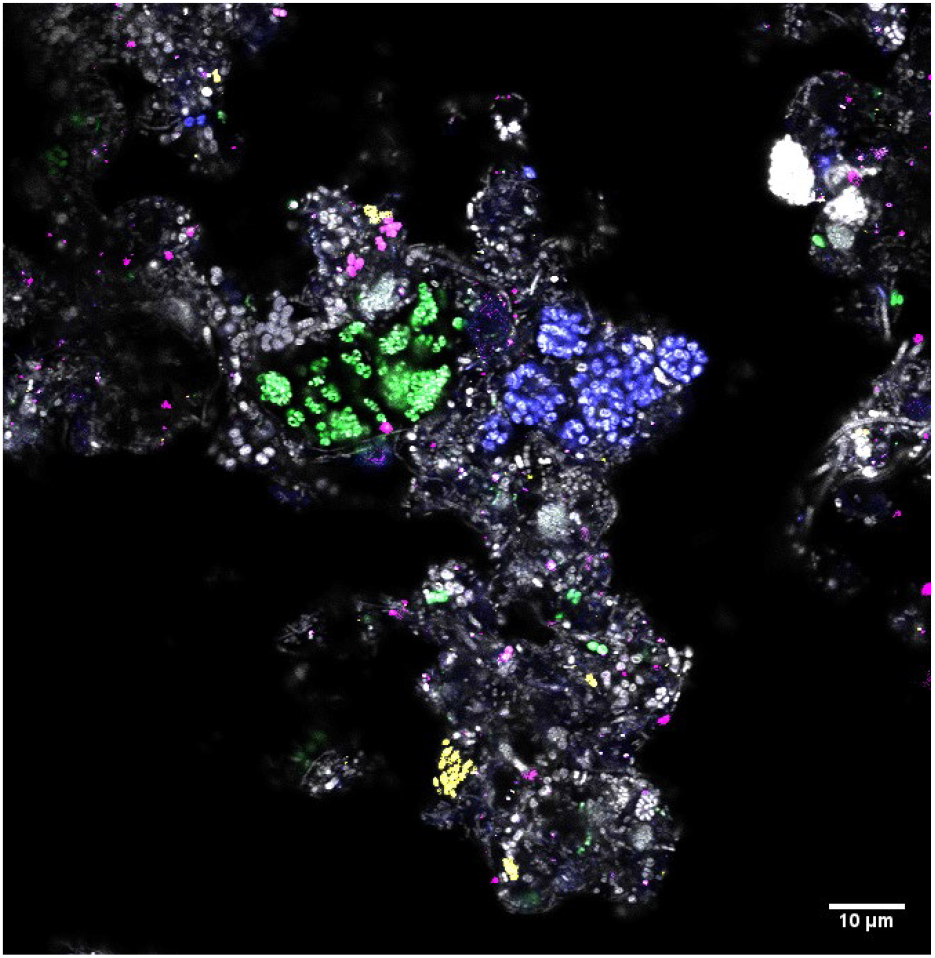
Multicolor FISH micrograph of different *Ca*. Accumulibacter species in full-scale activated sludge. *Ca*. A. proximus (green) was targeted by Acc471 probe. *Ca*. A. propinquus (blue) was targeted by Acc1011 probe. *Ca*. A. regalis (magenta) was targeted by Acc635 probe. *Ca*. A. delfensis and aalborgensis (yellow) were targeted by Acc470 probe. All bacteria (grey) were targeted with EUBmix probe. Scale bar is 10 µm.

The specificity of the widely applied PAOmix probe set (4) was evaluated *in silico* (**Figure 2**) and *in situ* (**Figure S6**). It showed lower specificity than expected, targeting various *Propionivibrio* spp., including *Ca*. Propionivibrio dominans *in situ*. Similarly, we tested *in silico* (**Figure 2**) the coverage and specificity of the “type” FISH probes Acc-I-444 and Acc-II-444 (19). While the Acc-I-444 targets several 16S rRNA gene sequences belonging to the *Ca*. A. aalborgensis and *Ca*. P. aalborgensis, Acc-II-444 showed a more specific coverage of the MiDAS-defined *Ca*. A. phosphatis cluster. The unspecific binding of the PAOmix and the “type” probes could explain why previous studies observed two populations of *Ca*. Accumulibacter with different morphologies (coccoid and rod-shaped) and wrongly concluded that there was a morphological difference between the two *ppk1*-defined types (16, 19).

### Potential for poly-P, glycogen and PHA accumulation and in situ validation of *Ca.* Accumulibacter and other related species

Genome mining for genes and pathways related to the PAO metabolism of the MAGs based on KEGG orthology was performed to confirm the potential for PAO metabolism. All the *Ca*. Accumulibacter spp. genomes encoded essential genes for polyphosphate accumulation and storage, such as the low-affinity phosphate transporter (*pit*) and the high affinity phosphate transporter (*pstSCAB*) (**Figure 5, Data S2**). The MAGs also encoded full potential for glycogen and PHA accumulation, typical of the PAO phenotype (**Figure 5, Data S2**). These metabolic predictions were further confirmed *in situ* by the presence of intracellular poly-P, PHA, and glycogen by FISH-Raman (**Table 2, Figure S7**). The MAGs belonging to *Ca*. Propionivibrio dominans encoded the full potential for PHA and glycogen accumulation, but not poly-P storage, as previously observed for *Ca*. P. aalborgensis (**Figure 5**, and (23)). These metabolic predictions were also confirmed by FISH-Raman analysis (**Figure S7**), which showed a similar intracellular profile for the *de novo* species midas_s_3472, representing additional evidence to support the different taxonomic classification of these lineages. Similarly, the MAGs belonging to *Ca*. Proximibacter danicus encoded the potential for poly-P and PHA accumulation, but only the latter was detected *in situ*.

**Figure 5.**
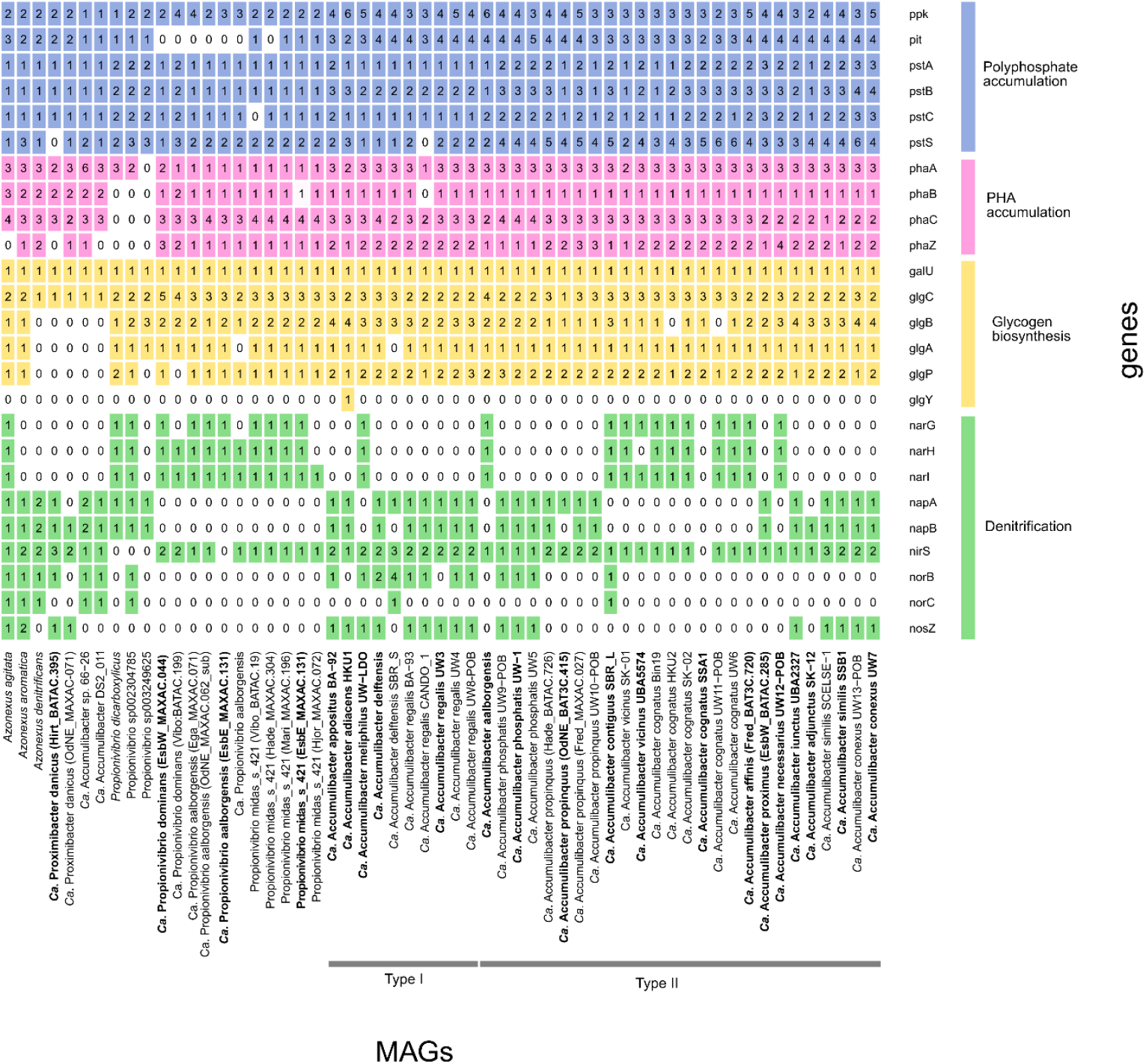
PAO metabolism-related functional potential of the *Ca*. Accumulibacter MAGs and closest relatives. The gene list follows the progression in the text. For the full list of gene names and associated KO numbers see Data S2. The MAGs and genomes are ordered as in the genome tree in Figure 1.

Differences in nitrate and nitrite reduction potential have often been suggested as a determining factor for niche and type differentiation, representing for a long time one of the most controversial (and arguably consequential) features of the *Ca*. Accumulibacter physiology (7, 19, 66, 67). Therefore, we also analyzed the distribution of genes involved in the denitrification process in *Ca*. Accumulibacter and related species (**Figure 5**). None of the *Ca*. Accumulibacter MAGs encoded the potential for full denitrification (nitrate to nitrogen gas). Genes encoding the respiratory nitrate reductase NarGHI were detected in *Ca*. Accumulibacter meliphilus (1 MAG), *Ca*. A. aalborgensis (1 MAG), *Ca*. Accumulibacter contiguus (1 MAG), *Ca*. Accumulibacter vicinus (2MAGs), *Ca*. Accumulibacter cognatus (5/6 MAGs), *Ca*. A. affinis (1 MAG) and Ca. Accumulibacter necessarius (1 MAG), while the other *Ca*. Accumulibacter MAGs encoded the genes for the periplasmic nitrate reductase NapAB. Although both enzymes can carry out nitrate reduction, the presence of the NarGHI enzyme complex was previously shown to be essential for anoxic phosphorus uptake using nitrate (66, 67). Its absence in the majority of the MAGs may indicate an inability to generate sufficient proton motive force to support respiration coupled to phosphorus uptake. On the contrary, the potential for nitrite reduction was more widespread, with *nirS* identified in all *Ca*. Accumulibacter MAGs, except *Ca*. A. cognatus. Nitric oxide reductase (*norBC*) was present only in 2 MAGs, representing *Ca*. A. deltensis SBR_S and *Ca*. A. contiguous SBR_L, while nitrous oxide reductase (*nosZ*) was encoded by *Ca*. A. appositus (1 MAG), *Ca*. A. adiacens (1 MAG), *Ca*. A. meliphilus (1 MAG), *Ca*. A. delftensis (1/2 MAGs), *Ca*. A. regalis (5/6 MAGs), *Ca*. A. iunctus (1 MAG), *Ca*. A. similis (2/2 MAG) and *Ca*. A. conexus (2/2 MAGs). Our metabolic predictions for *Ca*. A. delftensis slightly differed from Rubio-Rincon et al., (67), where they identified genes encoding the periplasmic nitrate reductase (*nap*) and a full set of nitrite (*nir*), nitric oxide (*nor*) and nitrous oxide (*nos*) reductases. Manual inspection of the *Ca*. A. delftensis genome in the MicroScope platform (48) revealed the presence of genes encoding for full reduction of nitrite to nitrogen gas. The distribution of the genes analysed in this study did not show any evidence supporting the hypothesis of a physiological distinction between the *pkk1*-defined type I and type II within the genus *Ca*. Accumulibacter. On the contrary, the specific set of genes involved in denitrification seems to be species-dependent.

Similarly, *Ca*. Propionivibrio dominans and *Ca*. Proximibacter danicus encoded genes for nitrate and nitrite reduction, differing as well in the nitrate reductase encoded (*narGHI* versus *napAB*, respectively). The *nirS* gene was identified in both. *Ca*. P. danicus MAGs, which encoded also nitrous oxide reductase (NosZ), which was missing in *Ca*. P. dominans. These metabolic differences across the genus *Ca*. Accumulibacter and the other related genera/species could explain why these different taxa could coexist in the same ecosystems and contribute to the overall phosphorus and nitrogen removal. However, experimental validation of these metabolic predictions, now achievable through the application of the species-specific FISH probes designed in this study in combination with microautoradiography (MAR-FISH) or Raman microspectroscopy, is needed to confirm metabolic abilities of these lineages and to determine their contribution to the full-scale EBPR process.

### Conclusions and perspectives

In this study, we provide a long-needed reassessment of the phylogeny of the genus *Ca*. Accumulibacter, using a comparison of genome-, *ppk1* and 16S rRNA gene approaches and identifying 18 novel species, for which we propose *Candidatus* names. We verified that the 16S rRNA gene is not able to resolve the phylogeny of these lineages and should be applied with caution in amplicon studies, as the results may be misleading. The *ppk1* gene is confirmed as the best choice for this purpose and offers a much higher resolution in distinguishing the different species. However, despite being 16S rRNA gene-based, a global survey such as the MiDAS4 can offer valuable insights to investigate the geographical distribution and major drivers of environmental filtering. As expected, *Ca*. Accumulibacter had a higher relative abundance in WWTPs performing biological phosphorus removal, indicating the process design as a major factor influencing their abundance. We also investigated the influence of the primer set chosen for the amplicon analysis and showed that despite being incapable of distinguish all the different species, the V1-V3 primer set was more suitable, compared to the V4 set which was unable to provide species-level resolution.

Finally, the species-specific FISH probes designed in this study, applied in combination with Raman microspectroscopy, confirmed the presence of the typical PAO storage polymers, predicted by metabolic annotation of the MAGs. Similarly, the MAGs were investigated for the distribution of genes encoding for the denitrification pathway related to one of the most controversial physiological traits of the *Ca*. Accumulibacter clades. None of the novel species encoded genes for full denitrification, but the annotation revealed fine-scale differences in the stepwise nitrogen-species reduction pathways, giving some insights into the niche differentiation of these lineages. Future experiments, for example using transcriptomics or activity-based studies with stable isotope-labelled compounds and FISH-Raman, could help to confirm the metabolic abilities of the different species and to explain how they could coexist in the same ecosystem and contribute to the overall phosphorus and nitrogen removal.

### Etymology

Proposed etymologies and protologues for the novel proposed species are provided in Tables S3-S19.

## Acknowledgements

We would like to thank Prof. Aharon Oren for his assistance with the naming etymology. The project was funded by the Villum Foundation (Dark Matter grant13351) and the United States National Science Foundation (MCB-1518130 to K.D.M.). The LABGeM (CEA/Genoscope & CNRS UMR8030), the France Génomique and French Bioinformatics Institute national infrastructures (funded as part of Investissement d’Avenir program managed by Agence Nationale pour la Recherche, contracts ANR-10-INBS-09 and ANR-11-INBS-0013) are acknowledged for support within the MicroScope annotation platform.

