## Supplementary Material for "Re-evaluation of the phylogenetic diversity and global distribution of the genus *Candidatus* Accumulibacter"

**Content:**

**Figure S1**. **Pairwise Genome-Wide Average Nucleotide Identity (ANI) Comparisons of *Ca.* Accumulibacter References.**

**Figure S2. Mean read abundance of 10 most abundant ASVs of different 16S- defined *Ca.* Accumulibacter species in global EBPR WWTPs, sorted by countries.**

**Figure S3. Global average relative read abundance of top 5 MiDAS-defined *Propionivibrio* species.**

**Figure S4. Global average relative read abundance of top 5 MiDAS-defined *Ca.* Proximibacter species**

**Figure S5. Evaluation of the V1-V3 and V4 primers for detection of *Ca*. Accummulibacter species.**

**Figure S6. Overlap of species-specific FISH probes for *Ca.* Accumulibacter and the widely applied probe PAO651**

**Table S1. Summary table of the FISH probes used in this study.**

**Table S2. Comparison of qFISH and 16S rRNA amplicon results (V1-V3).**

**Table S3. Protologue Table for *Candidatus* Accumulibacter regalis**

**Table S4. Protologue Table for *Candidatus* Accumulibacter appositus**

**Table S5. Protologue Table for *Candidatus* Accumulibacter adiacens**

**Table S6. Protologue Table for *Candidatus* Accumulibacter meliphilus**

**Table S7. Protologue Table for *Candidatus* Accumulibacter propinquus**

**Table S8. Protologue Table for *Candidatus* Accumulibacter contiguus**

**Table S9. Protologue Table for *Candidatus* Accumulibacter vicinus**

**Table S10. Protologue Table for *Candidatus* Accumulibacter cognatus**

**Table S11. Protologue Table for *Candidatus* Accumulibacter affinis**

**Table S12. Protologue Table for *Candidatus* Accumulibacter proximus**

**Table S13. Protologue Table for *Candidatus* Accumulibacter necessarius**

**Table S14. Protologue Table for *Candidatus* Accumulibacter iunctus**

**Table S15. Protologue Table for *Candidatus* Accumulibacter adjunctus**

**Table S16. Protologue Table for *Candidatus* Accumulibacter similis**

**Table S17. Protologue Table for *Candidatus* Accumulibacter conexus**

**Table S18. Protologue Table for *Candidatus* Propionivibrio dominans**

**Table S19. Protologue Table for *Candidatus* Proximibacter danicus**

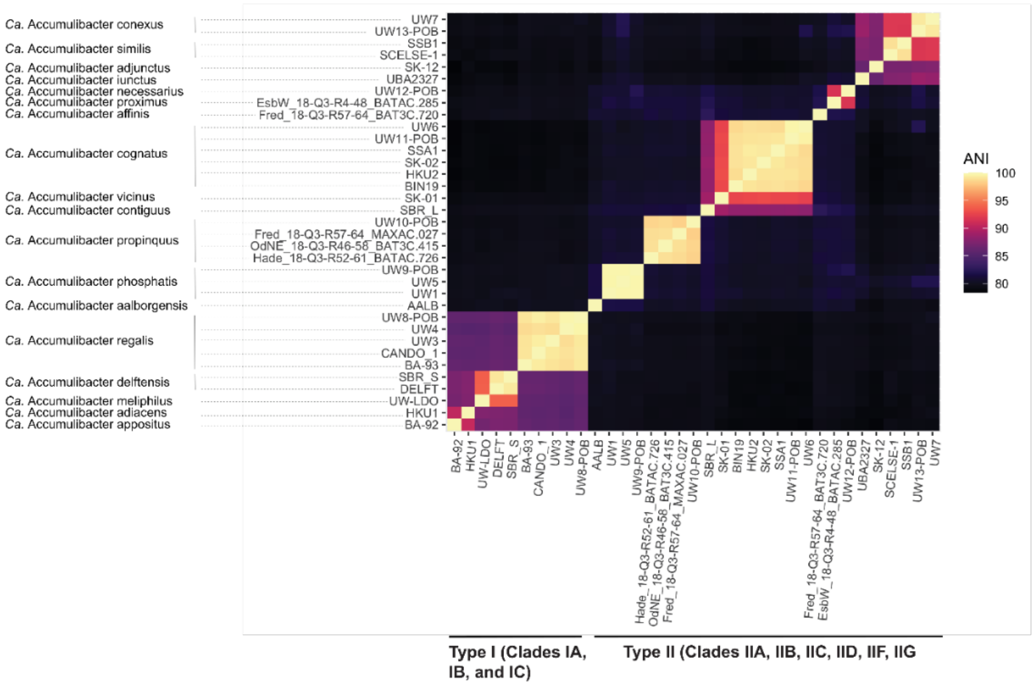


**Figure S1**. **Pairwise genome-wide Average Nucleotide Identity (ANI) comparisons of *Ca.* Accumulibacter references.** Pairwise ANI was calculated for all *Ca.* Accumulibacter genomes using fastANI. Each box in the heatmap represents the % ANI similarity between two MAGs, where MAGs are ordered by the genome tree in Figure 1. Proposed species names for representatives are provided on the y-axis and type and clade designations on the x-axis.

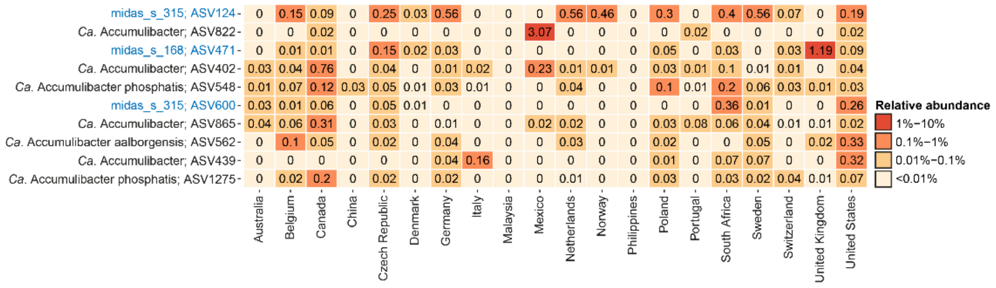


**Figure S2. The ten most abundant ASVs classified as *Ca.* Accumulibacter in EBPR plants worldwide.** Data represents the mean relative read abundance in EBPR plants across different countries and originates from the global survey of microbial communities in WWTPs (Dueholm et al., 2021), and it is based on V1-V3 amplicon dataset. Species marked in blue were wrongly classified as *Ca.* Accumulibacter according to the genome-based taxonomy.

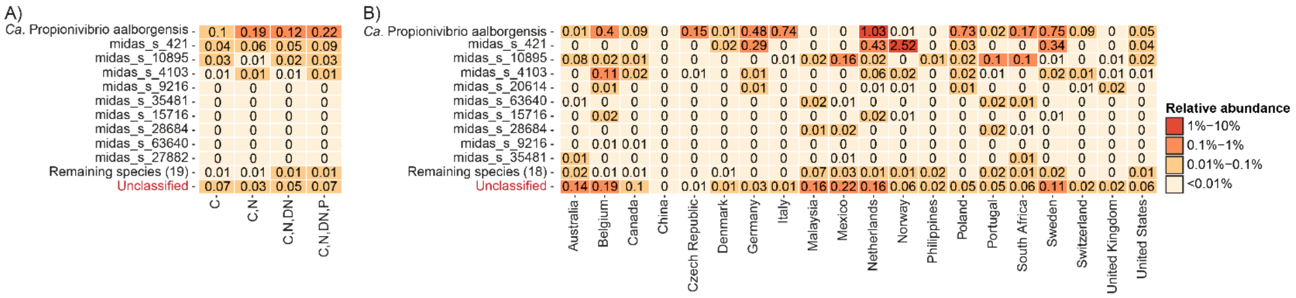


**Figure S3. The ten most abundant *Propionivibrio* species worldwide.** (A) Mean relative abundance across different process configurations (C - carbon removal; N – nitrification; DN - denitrification; P - phosphorus removal). (B) Mean relative abundance in EBPR plants across different across different countries. Data originates from the global survey of microbial communities in WWTPs (Dueholm et al., 2021), and it is based on V1-V3 amplicon dataset.

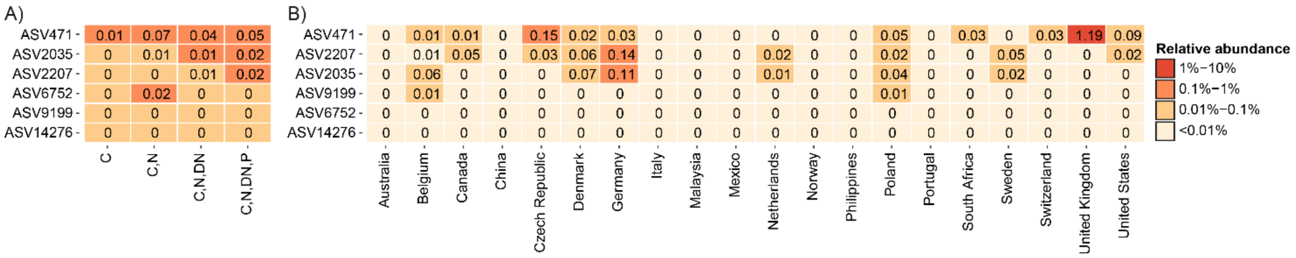


**Figure S4. ASVs classified as *Ca.* Proximibacter danicus (midas_s_168) worldwide.** (A) Mean relative abundance across different process configurations (C - carbon removal; N – nitrification; DN - denitrification; P - phosphorus removal). (B) Mean relative abundance in EBPR plants across different across different countries. Data originates from the global survey of microbial communities in WWTPs (Dueholm et al., 2021), and it is based on V1-V3 amplicon dataset.

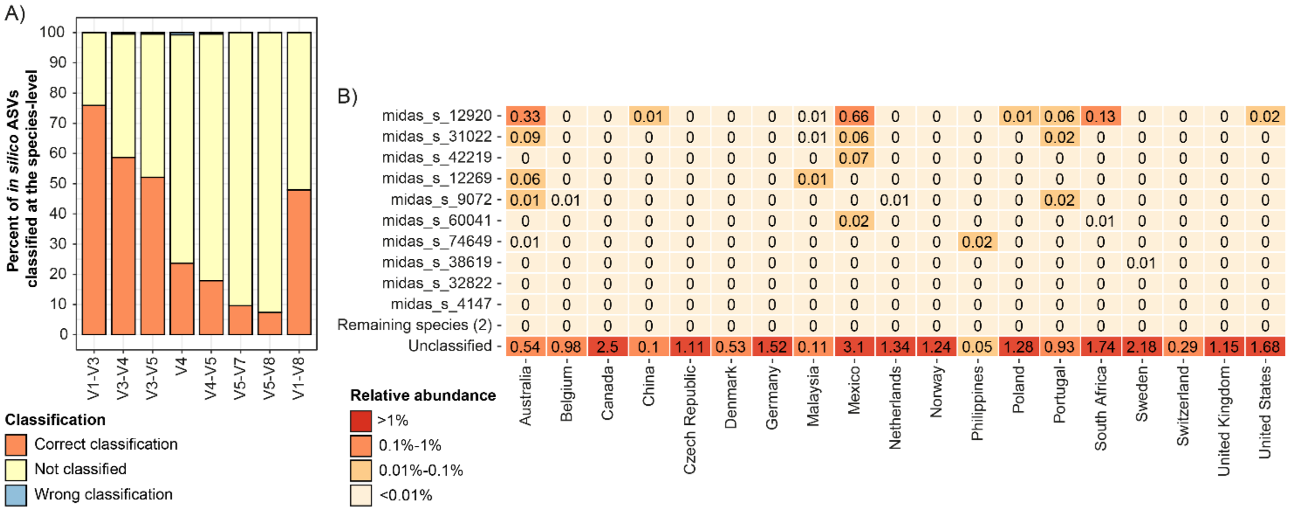


**Figure S5. Taxonomic resolution provided by different 16S rRNA amplicons and evaluation of V4 amplicons for detection of Ca. Accumulibacter species.** (A) Theoretical taxonomic signal in different variable regions of the 16S rRNA gene. *In silico* ASVs corresponding to different variable regions were extracted from all *Ca.* Accumulibacter reference sequences in the MiDAS 4.8.1 database. These short-reads were hereafter classified using the complete database. Classifications were evaluated at the species-level as correctly classified, wrongly classified, or not classified using the taxonomy of corresponding references as the ground truth. (B) Mean relative abundance of the ten most abundant *Ca.* Accumulibacter species in EBPR plants worldwide. Data originates from the global survey of microbial communities in WWTPs (Dueholm et al., 2021), and it is based on V4 amplicon dataset.

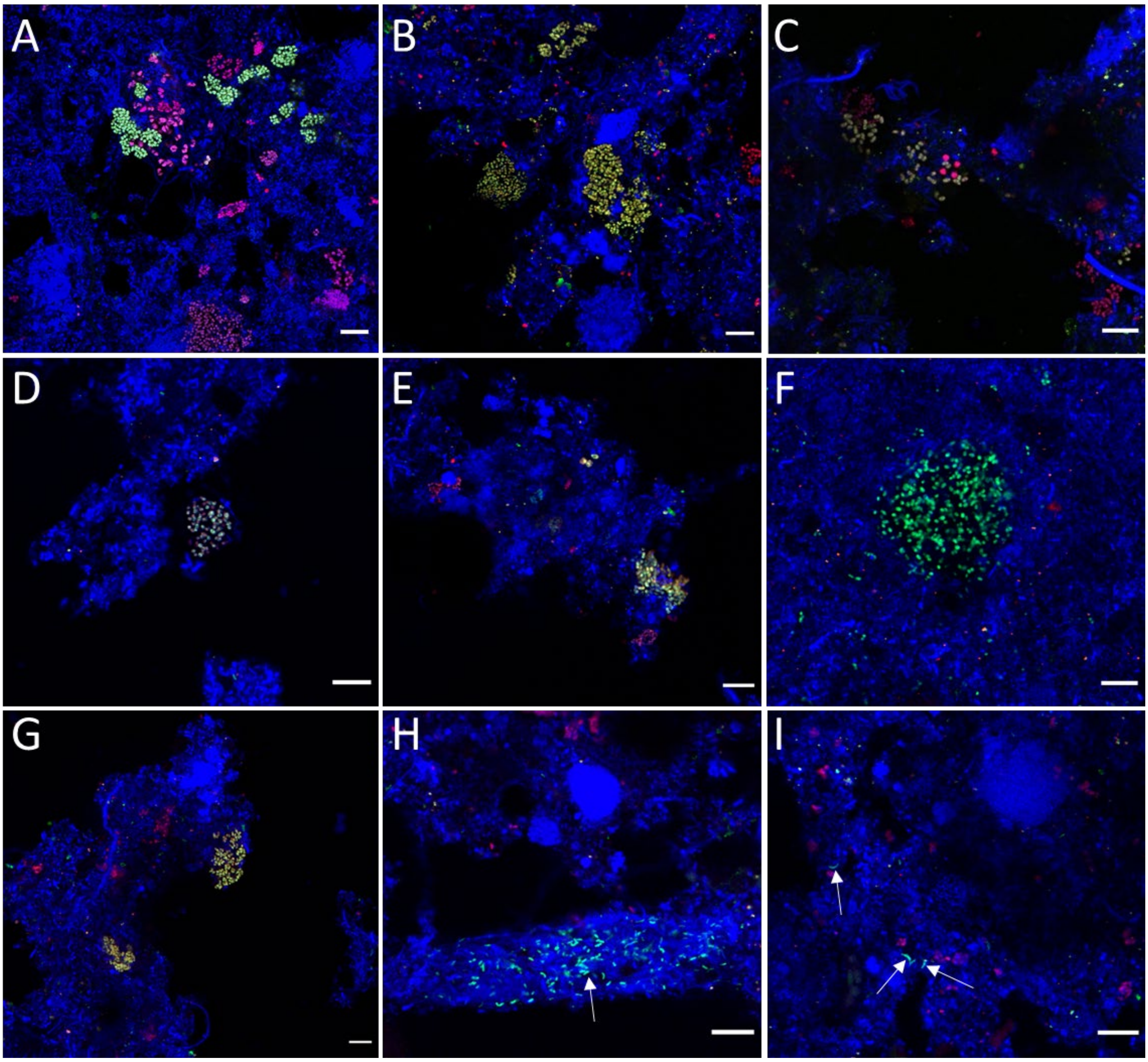


**Figure S6. Overlap of species-specific FISH probes for *Ca.* Accumulibacter and the widely applied probe PAO651.** (A) *Ca.* A. proximus (yellow-Acc469); (B) *Ca.* A. affinis and *Ca.* A. proximus (yellow-Acc471); (C) *Ca.* A. propinquus (yellow - Acc1011); (D) *Ca.* A. regalis (yellow - Acc635); (E) *Ca.* A. delfensis and *Ca.* A. aalborgensis (yellow - Acc470); (F) *Ca.* A. iunctus and *Ca.* A. similis (green - Acc741_2); (G) *Ca.* P. dominans (yellow - Acc213); (H) *Ca.* P. danicus (cyan - Acc442); (I) *Propionivibrio* sp. (cyan - midas_s_3472) (Acc441). In all pictures: EUBmix – blue, PAO651 – red, specific probe – green. Scale bar is indicating 10 µm.

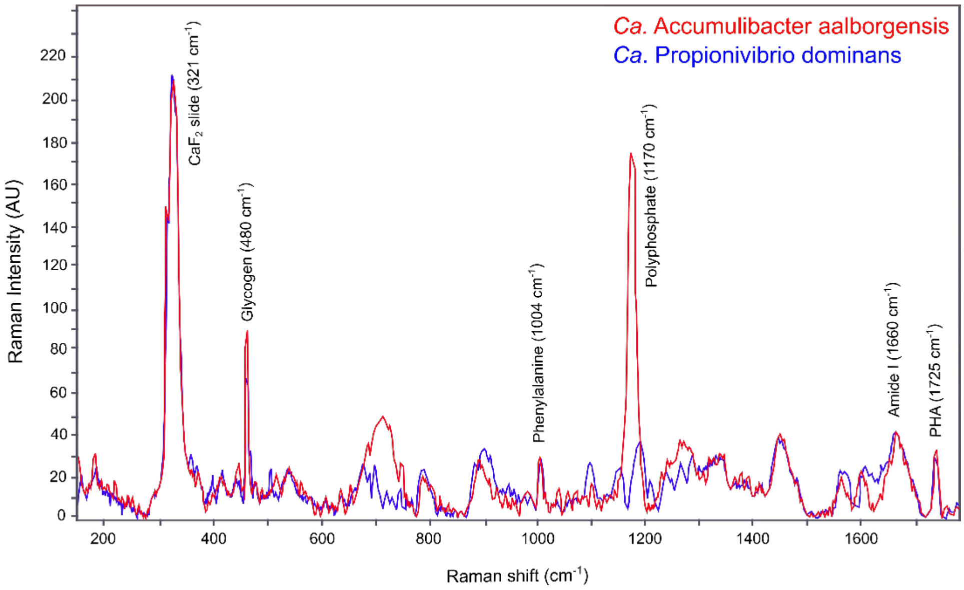


**Figure S7. Raman spectra of *Ca*. Accumulibacter aalborgensis and *Ca*. Propionivibrio dominans highlighting the differences in storage polymers content in the two genera.** Peaks for phenylalanine (1004 $cm^{-1}$) and amide I linkages of proteins (1660 $cm^{-1}$) are specific markers for biological material. Peaks for glycogen (480 $cm^{-1}$), polyphosphate (1170 $cm^{-1}$) and PHA (1725 $cm^{-1}$) are indicating the presence of the PAO-specific storage polymers. Spectra are calculated as average of 100 FISH-defined cells. AU, arbitrary units.

**Table S1. Summary table of the FISH probes used in this study.**

| Probe | *E. coli* pos. | Target group | Coverage * | Non-target hits | Sequence (5'-3') | [FA]%** | Reference |
|---|---|---|---|---|---|---|---|
| **PAO651** | **651–668** | **PAO cluster** | **44/467** | | **CCC TCT GCC AAA CTC CAG** | **35** | **(1)** |
| **Acc469** | **469-493** | ***Ca.* Accumulibacter proximus** | **2/3** | **0** | **CCA GGT ACC GTC ATC TAC ACA GGC** | **30** | **This study** |
| Acc469_C1 | 469-493 | Competitor for Acc469 | N/A | N/A | CCA GGT ACC GTC ATC TAC ACA GGG | N/A | This study |
| Acc469_C2 | 469-493 | Competitor for Acc469 | N/A | N/A | CTA GGT ACC GTC ATC TAC ACA GGC | N/A | This study |
| Acc469_C3 | 469-493 | Competitor for Acc469 | N/A | N/A | CWA GGT ACC GTC ATC TAC ACA GGG | N/A | This study |
| Acc469_C4 | 469-493 | Competitor for Acc469 | N/A | N/A | TCA GGT ACC GTC ATC TAC ACA GGG | N/A | This study |
| **Acc471** | **471-495** | ***Ca.* Accumulibacter affinis and proximus** | **39/61** | **1** | **CTC CAG GTA CCG TCA TCT ACA CAG** | **40** | **This study** |
| Acc471_C1 | 471-495 | Competitor for Acc471 | N/A | N/A | CTC CGG GTA CCG TCA TCT ACA CAG | N/A | This study |
| Acc471_C2 | 471-495 | Competitor for Acc471 | N/A | N/A | AGT CGG GTA CCG TCA TCT ACA CAG | N/A | This study |
| **Acc1011** | **1011-1032** | ***Ca.* Accumulibacter propinquus** | **8/61** | **3** | **GCG AGC ACT CCC AGA TCT CTC** | **40** | **This study** |
| Acc1011_C1 | 1011-1032 | Competitor for Acc1011 | N/A | N/A | GCG AGC ACT CCC AAA TCT CTC | N/A | This study |
| Acc1011_C2 | 1011-1032 | Competitor for Acc1011 | N/A | N/A | TCG AGC ACT CCC AGA TCT CTC | N/A | This study |
| Acc1011_C3 | 1011-1032 | Competitor for Acc1011 | N/A | N/A | GCG GGC ACT CCC AGA TCT CTC | N/A | This study |
| **Acc635** | **635-659** | ***Ca.* Accumulibacter regalis** | **11/61** | **0** | **AAC TCC AGC CTG GCA GTC TCA AAT** | **30** | **This study** |
| Acc635_C1 | 635-659 | Competitor for Acc635 | N/A | N/A | CAC TCC AGC CTG GCA GTC TCA AAT | N/A | This study |
| Acc635_C2 | 635-659 | Competitor for Acc635 | N/A | N/A | AAC TCC AGC CRG GCA GTC TCA AAT | N/A | This study |
| Acc635_C3 | 635-659 | Competitor for Acc635 | N/A | N/A | AAC TCC AGC TTG GCA GTC TCA AAT | N/A | This study |
| **Acc470** | **470-494** | ***Ca.* Accumulibacter aalborgensis and delftensis** | **68/86** | **0** | **TTC GGG TAC CGT CAT CTA CTC AGG** | **30** | **This study** |
| Acc470_C1 | 470-494 | Competitor for Acc470 | N/A | N/A | TGC GGG TAC CGT CAT CTA CTC AGG | N/A | This study |
| Acc470_C2 | 470-494 | Competitor for Acc470 | N/A | N/A | TTC GGG TAC CGT CAT CTA CAC AGG | N/A | This study |

| | | | | | | | |
|---|---|---|---|---|---|---|---|
| Acc470_C3 | 470-494 | Competitor for Acc470 | N/A | N/A | TTC GGG TAC CGT CAT CCA CTC AGA | N/A | This study |
| **Acc471_2** | **471-495** | ***Ca.* Accumulibacter iunctus and similis** | **18/21** | **3** | **AGT CGG GTA CCG TCA TCT ACA CAG** | **30** | **This study** |
| Acc471_2_C1 | 471-495 | Competitor for Acc471_2 | N/A | N/A | ATT CGG GTA CCG TCA TCT ACA CAG | N/A | This study |
| Acc471_2_C2 | 471-495 | Competitor for Acc471_2 | N/A | N/A | AGT CGG GTA CCG TCA TCG ACA CAG | N/A | This study |
| **Acc213** | **213- 235** | ***Ca.* Propionivibrio dominans** | **28/32** | **3** | **GGC CGC TCC TAA AGC AAG AGG T** | **35** | **This study** |
| Acc213_C1 | 213- 235 | Competitor for Acc213 | N/A | N/A | GGC CGC TCC CAA AGC AAG AGG T | N/A | This study |
| Acc213_C2 | 213- 235 | Competitor for Acc213 | N/A | N/A | GGC CGC TCC TAA AGC AAC AGG T | N/A | This study |
| **Acc442** | **442-464** | ***Ca.* Proximibacter danicus** | **17/22** | **2** | **GGA GAT GCG ATT TCT TCC CCG C** | **35** | **This study** |
| Acc442_C1 | 442-464 | Competitor for Acc442 | N/A | N/A | GGA GAT GCG ATT TCT TCC CRG | N/A | This study |
| **Acc441** | **441-461** | **midas_s_3472** | **19/34** | **2** | **AGT GCG ATT TCT TCC CCG CC** | **40** | **This study** |
| Acc441_C1 | 441-461 | Competitor for Acc441 | N/A | N/A | AGT GCH ATT TCT TCC CCG CC | N/A | This study |
| Acc441_C2 | 441-461 | Competitor for Acc441 | N/A | N/A | AGT GCG CTT TCT TCC CCG CC | N/A | This study |
| Acc441_C3 | 441-461 | Competitor for Acc441 | N/A | N/A | KGT GCG ATT TCT TCC CCG CC | N/A | This study |
| Acc441_C4 | 441-461 | Competitor for Acc441 | N/A | N/A | AGT GCG ATT TCT TCC CGG CC | N/A | This study |

* Taxonomy and coverage of groups is defined as in the MiDAS4 database. Values given as group hits/ group totals; ** Recommended optimal formamide concentration for use in FISH hybridizations; N/A – not applicable.

**Table S2. Comparison of qFISH and 16S rRNA amplicon results (V1-V3).** NA: not applicable.

| WWTP | Sample date | Abundance (%) | |
|---|---|---|---|
| | | Sequencing | qFISH |
| ***Ca.* Accumulibacter proximus (Acc469)** | | | |
| Odense NE, Denmark | October 2013 | NA | 1.7 ± 0.7 |
| Hjørring, Denmark | August 2011 | NA | <0.5 |
| Aalborg East, Denmark | March 2018 | NA | 0.8 ± 0.5 |
| ***Ca.* Accumulibacter affinis and proximus (Acc471)** | | | |
| Odense NE, Denmark | October 2013 | NA | 1.7 ± 0.6 |
| Hjørring, Denmark | August 2011 | NA | <0.5 |
| Aalborg East, Denmark | March 2018 | NA | 1.1 ± 0.7 |
| ***Ca.* Accumulibacter propinquus (Acc1011)** | | | |
| Odense NE, Denmark | October 2013 | NA | <0.5 |
| Hjørring, Denmark | August 2011 | NA | 0.6 ± 0.3 |
| Aalborg East, Denmark | March 2018 | NA | <0.5 |
| ***Ca.* Accumulibacter phosphatis (Acc635)** | | | |
| Odense NE, Denmark | October 2013 | NA | 1.3 ± 0.5 |
| Hjørring, Denmark | August 2011 | NA | 1.4 ± 0.9 |
| Aalborg East, Denmark | March 2018 | NA | 1 ± 0.5 |
| ***Ca.* Accumulibacter aalborgenesis (Acc470)** | | | |
| Fredericia, Denmark | October 2013 | NA | 4.8 ± 1.6 |
| Esbjerg West, Denmark | August 2013 | NA | 1.1 ± 0.4 |
| Esbjerg East, Denmark | October 2016 | NA | <0.5 |
| ***Ca.* Accumulibacter iunctus and similis (Acc471_2)** | | | |
| Tel Aviv, Israel | March 2018 | 1 | <0.5 |
| Limassol, Cyprus | February 2018 | 1.3 | <0.5 |
| Garmmexwolde, Netherlands | April 2018 | 2.7 | <0.5 |
| ***Ca.* Propionivibrio dominans (Acc213)** | | | |
| Boeslum, Denmark | August 2015 | 2.4 | <0.5 |
| Hjørring, Denmark | November 2018 | 1.4 | <0.5 |
| Piaseczno, Poland | March, 2018 | 2.8 | <0.5 |
| ***Ca.* Proximibacter danicus (Acc442)** | | | |
| Ribe, Denmark | October 2015 | 0.9 | <0.5 |
| East Anglia, England | March 2018 | 0.8 | <0.5 |
| J-town Germany | March 2018 | 0.8 | <0.5 |
| **midas_s_3472 (Acc441)** | | | |
| Esbjerg W, Denmark | February 2017 | 2 | <0.5 |
| Viborg, Denmark | February 2013 | 0.8 | <0.5 |
| Garmmexwolde, Netherlands | April 2018 | 2.7 | <0.5 |

**Table S3. Protologue Table for *Candidatus* Accumulibacter regalis**

| | |
|---|---|
| Species name | *Candidatus* Accumulibacter regalis |
| Genus name | *Candidatus* Accumulibacter |
| Specific epithet | regalis |
| Type species of the genus | *Candidatus* Accumulibacter phosphatis |
| Genus status | Candidatus |
| Species etymology | Description of "*Candidatus* Accumulibacter regalis" sp. nov.: "*Candidatus* Accumulibacter regalis," (re.ga´lis, L. masc. adj. regalis, royal; indicating the dominating role in the bioreactors from which the MAG was retrieved). |
| Species status | sp. nov. |
| Designation of the type MAG | GCF_000024165.1 |
| MAG/SAG accession number | IMG accession ID 2687453699 |
| Genome status | High-quality draft |
| Genome size | 4329894 |
| GC mol % | 63.9 |
| Country of origin | USA |
| Region of origin | Wisconsin |
| Source of sample | EBPR bioreactor |
| Geographical location | Madison |
| Latitude | - |
| Longitude | - |
| Depth | - |
| Altitude | - |
| Temperature of the sample | - |
| pH of the sample | - |
| Relationship to oxygen | Facultative anaerobic |
| Energy metabolism | Polyphosphate-accumulating organism |
| Assembly | - |
| Sequencing technology | - |
| Binning software used | - |
| Assembly software used | - |
| Habitat | - |
| Miscellaneous, extraordinary features relevant for the description | - |

**Table S4. Protologue Table for *Candidatus* Accumulibacter appositus**

| | |
|---|---|
| Species name | *Candidatus* Accumulibacter appositus |
| Genus name | *Candidatus* Accumulibacter |
| Specific epithet | appositus |
| Type species of the genus | *Candidatus* Accumulibacter phosphatis |
| Genus status | Candidatus |
| Species etymology | Description of "*Candidatus* Accumulibacter appositus" sp. nov.: "*Candidatus* Accumulibacter appositus," (ap.po´si.tus, L. masc. part. adj. *appositus*, close; indicating the close phylogenetic relationship with *Ca*. Accumulibacter phosphatis). This taxon is represented by the MAG *Candidatus* Accumulibacter sp. BA-92. |
| Species status | sp. nov. |
| Designation of the type MAG | GCA_000585055.1 |
| MAG/SAG accession number | GCF_000024165.1 |
| Genome status | High-quality draft |
| Genome size | 4947936 |
| GC mol % | 62.93 |
| Country of origin | Australia |
| Region of origin | - |
| Source of sample | Enriched from a lab-scale EBPR bioreactor |
| Geographical location | - |
| Latitude | 27.4860 |
| Longitude | 153.1907 |
| Depth | - |
| Altitude | - |
| Temperature of the sample | - |
| pH of the sample | - |
| Relationship to oxygen | Facultative anaerobic |
| Energy metabolism | Polyphosphate-accumulating organism |
| Assembly | 1 sample/Scaffold – not sure about this |
| Sequencing technology | Illumina HiSeq |
| Binning software used | - |
| Assembly software used | CLC Genomics Workbench v. 5.5 |
| Habitat | Enriched from a lab-scale EBPR bioreactor |
| Miscellaneous, extraordinary features relevant for the description | - |

**Table S5. Protologue Table for *Candidatus* Accumulibacter adiacens**

| | |
|---|---|
| Species name | *Candidatus* Accumulibacter adiacens |
| Genus name | *Candidatus* Accumulibacter |
| Specific epithet | adiacens |
| Type species of the genus | *Candidatus* Accumulibacter phosphatis |
| Genus status | Candidatus |
| Species etymology | Description of "*Candidatus* Accumulibacter adiacens" sp. nov.: "*Candidatus* Accumulibacter adiacens," (ad'ia.cens, L. part. adj. *adiacens*, close; indicating the close phylogenetic relationship with *Ca*. Accumulibacter phosphatis). This taxon is represented by the MAG *Candidatus* Accumulibacter phosphatis HKU1. |
| Species status | sp. nov. |
| Designation of the type MAG | GCF_000024165.1 |
| MAG/SAG accession number | GCA_000987445.1 |
| Genome status | High-quality draft |
| Genome size | 3625256 |
| GC mol % | 63.89 |
| Country of origin | Hong Kong |
| Region of origin | - |
| Source of sample | bioreactor treating saline wastewater (1% salinity) containing high content of P |
| Geographical location | - |
| Latitude | 22.28 |
| Longitude | 114.14 |
| Depth | N/A |
| Altitude | N/A |
| Temperature of the sample | - |
| pH of the sample | N/A |
| Relationship to oxygen | Facultative anaerobe |
| Energy metabolism | Polyphosphate-accumulating organism |
| Assembly | 1 sample |
| Sequencing technology | Illumina HiSeq2000 |
| Binning software used | - |
| Assembly software used | CLC Genomics Workbench v. 6.0.2 |
| Habitat | bioreactor treating saline wastewater (1% salinity) containing high content of P |
| Miscellaneous, extraordinary features relevant for the description | - |

**Table S6. Protologue Table for *Candidatus* Accumulibacter meliphilus**

| | |
|---|---|
| Species name | *Candidatus* Accumulibacter meliphilus |
| Genus name | *Candidatus* Accumulibacter |
| Specific epithet | meliphilus |
| Type species of the genus | *Candidatus* Accumulibacter phosphatis |
| Genus status | Candidatus |
| Species etymology | Description of "*Candidatus* Accumulibacter meliphilus" sp. nov.: "*Candidatus* Accumulibacter meliphilus," (me.li'phi.lus. L. fem. n. *meles*, badger; Gr. masc. n. *philos*, friend; N.L. masc. n. *meliphilus*, friend of the badger, indicating the badger mascot of the University of Wisconsin-Madison, where the MAG was retrieved). This taxon is represented by the MAG *Candidatus* Accumulibacter UW-LDO. |
| Species status | sp. nov. |
| Designation of the type MAG | GCF_000024165.1 |
| MAG/SAG accession number | GCA_003332265.1 |
| Genome status | High-quality draft |
| Genome size | 4,701,482 |
| GC mol % | - |
| Country of origin | USA |
| Region of origin | Wisconsin |
| Source of sample | - |
| Geographical location | Madison |
| Latitude | - |
| Longitude | - |
| Depth | - |
| Altitude | - |
| Temperature of the sample | - |
| pH of the sample | - |
| Relationship to oxygen | Facultative anaerobe |
| Energy metabolism | Polyphosphate-accumulating organism |
| Assembly | - |
| Sequencing technology | Illumina HiSeq; Nanopore |
| Binning software used | - |
| Assembly software used | SPAdes v. 3.9.0; Anvi'o v. 4; Links v. 1.5 |
| Habitat | - |
| Miscellaneous, extraordinary features relevant for the description | - |

**Table S7. Protologue Table for *Candidatus* Accumulibacter propinquus**

| | |
|---|---|
| Species name | *Candidatus* Accumulibacter propinquus |
| Genus name | *Candidatus* Accumulibacter |
| Specific epithet | propinquus |
| Type species of the genus | *Candidatus* Accumulibacter phosphatis |
| Genus status | Candidatus |
| Species etymology | Description of "*Candidatus* Accumulibacter propinquus" sp. nov.: "*Candidatus* Accumulibacter propinquus," (pro.pin´qu.us L. masc. adj. propinquus, next of kin; indicating the close phylogenetic relationship with *Ca*. Accumulibacter phosphatis). This taxon is represented by the MAG OdNE_BAT3C.415. |
| Species status | sp. nov. |
| Designation of the type MAG | GCF_000024165.1 |
| MAG/SAG accession number | GCA_016714935.1 |
| Genome status | High-quality draft |
| Genome size | 4325360 |
| GC mol % | 63.00 |
| Country of origin | Denmark |
| Region of origin | Odense East |
| Source of sample | Full-scale enriched biological phosphorus removal wastewater treatment plant |
| Geographical location | Odense East |
| Latitude | 55.432604 |
| Longitude | 10.45886 |
| Depth | - |
| Altitude | - |
| Temperature of the sample | Mesophilic |
| pH of the sample | - |
| Relationship to oxygen | Facultative anaerobe |
| Energy metabolism | Polyphosphate-accumulating organism |
| Assembly | 1 sample |
| Sequencing technology | Oxford Nanopore and Illumina Hiseq X |
| Binning software used | MetaBAT2 |
| Assembly software used | CANU v1.8 |
| Habitat | Full-scale enriched biological phosphorus removal wastewater treatment plant |
| Miscellaneous, extraordinary features relevant for the description | Coccoid-shaped cell with a diameter of approx. 0.8-1.2 µm. |

**Table S8. Protologue Table for *Candidatus* Accumulibacter contiguus**

| | |
|---|---|
| Species name | *Candidatus* Accumulibacter contiguus |
| Genus name | *Candidatus* Accumulibacter |
| Specific epithet | contiguus |
| Type species of the genus | *Candidatus* Accumulibacter phosphatis |
| Genus status | Candidatus |
| Species etymology | Description of "*Candidatus* Accumulibacter contiguus" sp. nov.: "*Candidatus* Accumulibacter contiguus," (con.ti´gu.us. L. masc. adj. *contiguus*, close; indicating the close phylogenetic relationship with *Ca*. Accumulibacter phosphatis). This taxon is represented by the MAG *Candidatus* Accumulibacter phosphatis SBR_L. |
| Species status | sp. nov. |
| Designation of the type MAG | GCF_000024165.1 |
| MAG/SAG accession number | GCA_012940005.1 |
| Genome status | High-quality draft |
| Genome size | 502444 |
| GC mol % | 61.73 |
| Country of origin | USA |
| Region of origin | Columbia |
| Source of sample | enrichment culture grown in lab-scale EBPR SBR; organism purified/enriched using density-based techniques |
| Geographical location | Columbia |
| Latitude | - |
| Longitude | - |
| Depth | - |
| Altitude | - |
| Temperature of the sample | - |
| pH of the sample | - |
| Relationship to oxygen | Facultative anaerobe |
| Energy metabolism | Polyphosphate-accumulating organism |
| Assembly | - |
| Sequencing technology | IonTorrent |
| Binning software used | - |
| Assembly software used | - |
| Habitat | enrichment culture grown in lab-scale EBPR SBR; organism purified/enriched using density-based techniques |
| Miscellaneous, extraordinary features relevant for the description | - |

**Table S9. Protologue Table for *Candidatus* Accumulibacter vicinus**

| | |
|---|---|
| Species name | *Candidatus* Accumulibacter vicinus |
| Genus name | *Candidatus* Accumulibacter |
| Specific epithet | vicinus |
| Type species of the genus | *Candidatus* Accumulibacter phosphatis |
| Genus status | Candidatus |
| Species etymology | Description of “*Candidatus* Accumulibacter vicinus” sp. nov.: “*Candidatus* Accumulibacter vicinus,” (vi.ci´nus. L. masc. adj. *vicinus*, close; indicating the close phylogenetic relationship with *Ca*. Accumulibacter phosphatis). This taxon is represented by the MAG Accumulibacter phosphatis UBA5574. |
| Species status | sp. nov. |
| Designation of the type MAG | GCF_000024165.1 |
| MAG/SAG accession number | GCA_002425405.1 |
| Genome status | High-quality draft |
| Genome size | 428324 |
| GC mol % | 61.98 |
| Country of origin | Australia |
| Region of origin | Queensland |
| Source of sample | - |
| Geographical location | Queensland |
| Latitude | - |
| Longitude | - |
| Depth | - |
| Altitude | - |
| Temperature of the sample | - |
| pH of the sample | - |
| Relationship to oxygen | Facultative anaerobe |
| Energy metabolism | Polyphosphate-accumulating organism |
| Assembly | - |
| Sequencing technology | Illumina |
| Binning software used | - |
| Assembly software used | CLC de novo assembler v. 4.4.1 |
| Habitat | - |
| Miscellaneous, extraordinary features relevant for the description | - |

**Table S10. Protologue Table for *Candidatus* Accumulibacter cognatus**

| | |
|---|---|
| Species name | *Candidatus* Accumulibacter cognatus |
| Genus name | *Candidatus* Accumulibacter |
| Specific epithet | cognatus |
| Type species of the genus | *Candidatus* Accumulibacter phosphatis |
| Genus status | Candidatus |
| Species etymology | Description of "*Candidatus* Accumulibacter cognatus" sp. nov.: "*Candidatus* Accumulibacter cognatus," (co.gna´tus, L. masc. adj. *cognatus*, similar; indicating the close phylogenetic relationship with *Ca*. Accumulibacter phosphatis). This taxon is represented by the MAG *Candidatus* Accumulibacter phosphatis SSA1. |
| Species status | sp. nov. |
| Designation of the type MAG | GCF_000024165.1 |
| MAG/SAG accession number | GCA_013414765.1 |
| Genome status | High-quality draft |
| Genome size | 516951 |
| GC mol % | 61.4 |
| Country of origin | Singapore |
| Region of origin | Singapore |
| Source of sample | Activated sludge, continuous culture bioreactor |
| Geographical location | Jurong West |
| Latitude | 1.344722 |
| Longitude | 103.681389 |
| Depth | - |
| Altitude | - |
| Temperature of the sample | - |
| pH of the sample | - |
| Relationship to oxygen | Facultative anaerobe |
| Energy metabolism | Polyphosphate-accumulating organism |
| Assembly | 1 sample |
| Sequencing technology | Oxford Nanopore MiniION |
| Binning software used | - |
| Assembly software used | Canu v. 1.8 |
| Habitat | Activated sludge, continuous culture bioreactor |
| Miscellaneous, extraordinary features relevant for the description | - |

**Table S11. Protologue Table for *Candidatus* Accumulibacter affinis**

| | |
|---|---|
| Species name | *Candidatus* Accumulibacter affinis |
| Genus name | *Candidatus* Accumulibacter |
| Specific epithet | affinis |
| Type species of the genus | *Candidatus* Accumulibacter phosphatis |
| Genus status | Candidatus |
| Species etymology | Description of "*Candidatus* Accumulibacter affinis" sp. nov.: "*Candidatus* Accumulibacter affinis," (af.fi´nis. L. masc. adj. affinis, next of kin; indicating the close phylogenetic relationship with *Ca*. Accumulibacter phosphatis). This taxon is represented by the MAG Fred_BAT3C.720. |
| Species status | sp. nov. |
| Designation of the type MAG | GCF_000024165.1 |
| MAG/SAG accession number | GCA_016713625.1 |
| Genome status | High-quality draft |
| Genome size | 5041483 |
| GC mol % | 62.30 |
| Country of origin | Denmark |
| Region of origin | Fredericia |
| Source of sample | Full-scale enriched biological phosphorus removal wastewater treatment plant |
| Geographical location | Fredericia |
| Latitude | 55.552368 |
| Longitude | 9.720404 |
| Depth | - |
| Altitude | - |
| Temperature of the sample | Mesophilic |
| pH of the sample | - |
| Relationship to oxygen | Facultative anaerobe |
| Energy metabolism | Polyphosphate-accumulating organism |
| Assembly | 1 sample |
| Sequencing technology | Oxford Nanopore and Illumina Hiseq X |
| Binning software used | MetaBAT2 |
| Assembly software used | CANU v1.8 |
| Habitat | Full-scale enriched biological phosphorus removal wastewater treatment plant |
| Miscellaneous, extraordinary features relevant for the description | Coccoid-shaped cells with a diameter approx. of 0.5-0.7 µm |

**Table S12. Protologue Table for *Candidatus* Accumulibacter proximus**

| | |
|---|---|
| Species name | *Candidatus* Accumulibacter proximus |
| Genus name | *Candidatus* Accumulibacter |
| Specific epithet | proximus |
| Type species of the genus | *Candidatus* Accumulibacter phosphatis |
| Genus status | Candidatus |
| Species etymology | Description of "*Candidatus* Accumulibacter proximus" sp. nov.: "*Candidatus* Accumulibacter proximus," (pro´xi.mus L. masc. adj. proximus, next of kin; indicating the close phylogenetic with *Ca*. Accumulibacter phosphatis). This taxon is represented by the MAG EsbW_BATAC.285. |
| Species status | sp. nov. |
| Designation of the type MAG | GCF_000024165.1 |
| MAG/SAG accession number | GCA_016709675.1 |
| Genome status | High-quality draft |
| Genome size | 5402764 |
| GC mol % | 62.70 |
| Country of origin | Denmark |
| Region of origin | Esbjerg West |
| Source of sample | Full-scale biological nutrient removal wastewater treatment plant |
| Geographical location | Esbjerg West |
| Latitude | 55.488097 |
| Longitude | 8.430505 |
| Depth | - |
| Altitude | - |
| Temperature of the sample | Mesophilic |
| pH of the sample | - |
| Relationship to oxygen | Facultative anaerobe |
| Energy metabolism | Polyphosphate-accumulating organism |
| Assembly | 1 sample |
| Sequencing technology | Oxford Nanopore and Illumina Hiseq X |
| Binning software used | MetaBAT2 |
| Assembly software used | CANU v1.8 |
| Habitat | Full-scale biological nutrient removal wastewater treatment plant |
| Miscellaneous, extraordinary features relevant for the description | Coccoid-shaped cells with a diameter approx. of 0.5-0.7 µm |

**Table S13. Protologue Table for *Candidatus* Accumulibacter necessarius**

| | |
|---|---|
| Species name | *Candidatus* Accumulibacter necessarius |
| Genus name | *Candidatus* Accumulibacter |
| Specific epithet | necessarius |
| Type species of the genus | *Candidatus* Accumulibacter phosphatis |
| Genus status | Candidatus |
| Species etymology | Description of "*Candidatus* Accumulibacter necessarius" sp. nov.: "*Candidatus* Accumulibacter necessarius," (ne.ces.sa´ri.us. L. masc. adj. *necessarius*, next of kin; indicating the close phylogenetic relationship with *Ca*. Accumulibacter phosphatis). This taxon is represented by the MAG *Candidatus* Accumulibacter UW12-POB. |
| Species status | sp. nov. |
| Designation of the type MAG | GCF_000024165.1 |
| MAG/SAG accession number | GCA_017302435.1 |
| Genome status | High-quality draft |
| Genome size | 456421 |
| GC mol % | 62.7 |
| Country of origin | USA |
| Region of origin | Wisconsin |
| Source of sample | Activated sludge |
| Geographical location | Madison |
| Latitude | - |
| Longitude | - |
| Depth | - |
| Altitude | - |
| Temperature of the sample | - |
| pH of the sample | - |
| Relationship to oxygen | Facultative anaerobe |
| Energy metabolism | Polyphosphate-accumulating organism |
| Assembly | 1 sample |
| Sequencing technology | Illumina HiSeq |
| Binning software used | - |
| Assembly software used | metaSPAdes v. 3.9.0 |
| Habitat | Activated sludge |
| Miscellaneous, extraordinary features relevant for the description | - |

**Table S14. Protologue Table for *Candidatus* Accumulibacter iunctus**

| | |
|---|---|
| Species name | *Candidatus* Accumulibacter iunctus |
| Genus name | *Candidatus* Accumulibacter |
| Specific epithet | iunctus |
| Type species of the genus | *Candidatus* Accumulibacter phosphatis |
| Genus status | Candidatus |
| Species etymology | Description of "*Candidatus* Accumulibacter iunctus" sp. nov.: "*Candidatus* Accumulibacter iunctus," (iunc´tus L. masc. adj. iunctus, next of kin; indicating the close phylogenetic relationship with *Ca*. Accumulibacter phosphatis). This taxon is represented by the MAG *Candidatus* Accumulibacter UBA2327. |
| Species status | sp. nov. |
| Designation of the type MAG | GCF_000024165.1 |
| MAG/SAG accession number | GCA 002345025.1 |
| Genome status | High-quality draft |
| Genome size | 4431027 |
| GC mol % | 65.20 |
| Country of origin | Australia |
| Region of origin | Queensland |
| Source of sample | Bioreactor sludge metagenome |
| Geographical location | Brisbane, Thorneside Wastewater Treatment Plant |
| Latitude | - |
| Longitude | - |
| Depth | - |
| Altitude | - |
| Temperature of the sample | - |
| pH of the sample | - |
| Relationship to oxygen | Facultative anaerobe |
| Energy metabolism | Polyphosphate-accumulating organism |
| Assembly | 1 sample |
| Sequencing technology | Illumina HiSeq |
| Binning software used | - |
| Assembly software used | CLC de novo assembler v. 4.4.1 |
| Habitat | Activated sludge |
| Miscellaneous, extraordinary features relevant for the description | - |

**Table S15. Protologue Table for *Candidatus* Accumulibacter adjunctus**

| | |
|---|---|
| Species name | *Candidatus* Accumulibacter adjunctus |
| Genus name | *Candidatus* Accumulibacter |
| Specific epithet | adiunctus |
| Type species of the genus | *Candidatus* Accumulibacter phosphatis |
| Genus status | Candidatus |
| Species etymology | Description of "Candidatus Accumulibacter adjunctus" sp. nov.: "Candidatus Accumulibacter adjunctus," (ad.iunc´tus. L. masc. part. adj. adjunctus, close; indicating the close phylogenetic relationship with Ca. Accumulibacter phosphatis). This taxon is represented by the MAG *Candidatus* Accumulibacter sp. SK-12 |
| Species status | sp. nov. |
| Designation of the type MAG | GCF_000024165.1 |
| MAG/SAG accession number | GCA_000585015.1 |
| Genome status | High-quality draft |
| Genome size | 4,412,715 |
| GC mol % | 65.80 |
| Country of origin | Australia |
| Region of origin | Queensland |
| Source of sample | synthetic wastewater |
| Geographical location | Brisbane, Thorneside Wastewater Treatment Plant |
| Latitude | 27.485973 |
| Longitude | 153.190699 |
| Depth | - |
| Altitude | - |
| Temperature of the sample | - |
| pH of the sample | - |
| Relationship to oxygen | Facultative anaerobe |
| Energy metabolism | Polyphosphate-accumulating organism |
| Assembly | - |
| Sequencing technology | Illumina HiSeq |
| Binning software used | - |
| Assembly software used | CLC Genomics Workbench v. 5.5 |
| Habitat | EBPR sludge |
| Miscellaneous, extraordinary features relevant for the description | - |

**Table S16. Protologue Table for *Candidatus* Accumulibacter similis**

| | |
|---|---|
| Species name | *Candidatus* Accumulibacter similis |
| Genus name | *Candidatus* Accumulibacter |
| Specific epithet | similis |
| Type species of the genus | *Candidatus* Accumulibacter phosphatis |
| Genus status | Candidatus |
| Species etymology | Description of "*Candidatus* Accumulibacter similis" sp. nov.: "*Candidatus* Accumulibacter similis," (si´mi.lis. L. masc. adj. *similis*, similar; indicating the close phylogenetic relationship with *Ca*. Accumulibacter phosphatis). This taxon is represented by the MAG *Candidatus* Accumulibacter phosphatis SSB1. |
| Species status | sp. nov. |
| Designation of the type MAG | GCF_000024165.1 |
| MAG/SAG accession number | GCA_013347225.1 |
| Genome status | High-quality draft |
| Genome size | 503901 |
| GC mol % | 66.0 |
| Country of origin | Singapore |
| Region of origin | Jurong West |
| Source of sample | Activated sludge |
| Geographical location | Jurong West |
| Latitude | 1.344722 |
| Longitude | 103.681389 |
| Depth | - |
| Altitude | - |
| Temperature of the sample | - |
| pH of the sample | - |
| Relationship to oxygen | Facultative anaerobe |
| Energy metabolism | Polyphosphate-accumulating organism |
| Assembly | 1 sample |
| Sequencing technology | Oxford Nanopore MiniION |
| Binning software used | - |
| Assembly software used | Canu v. 1.8 |
| Habitat | Activated sludge |
| Miscellaneous, extraordinary features relevant for the description | - |

**Table S17. Protologue Table for *Candidatus* Accumulibacter conexus**

| | |
|---|---|
| Species name | *Candidatus* Accumulibacter conexus |
| Genus name | *Candidatus* Accumulibacter |
| Specific epithet | conexus |
| Type species of the genus | *Candidatus* Accumulibacter phosphatis |
| Genus status | Candidatus |
| Species etymology | Description of "*Candidatus* Accumulibacter conexus" sp. nov.: "*Candidatus* Accumulibacter conexus," (co.ne´xus. L. masc. part. adj. *conexus*, next of kin; indicating the close phylogenetic relationship with *Ca*. Accumulibacter phosphatis). This taxon is represented by the MAG *Candidatus* Accumulibacter UW7 |
| Species status | sp. nov. |
| Designation of the type MAG | GCF_000024165.1 |
| MAG/SAG accession number | GCA_017592775.1 |
| Genome status | High-quality draft |
| Genome size | 487076 |
| GC mol % | 66.30 |
| Country of origin | USA |
| Region of origin | Wisconsin |
| Source of sample | Activate sludge |
| Geographical location | Wisconsin |
| Latitude | 43.071 |
| Longitude | 89.401 |
| Depth | - |
| Altitude | - |
| Temperature of the sample | - |
| pH of the sample | - |
| Relationship to oxygen | Facultative anaerobe |
| Energy metabolism | Polyphosphate-accumulating organism |
| Assembly | 1 sample |
| Sequencing technology | Illumina HiSeq |
| Binning software used | - |
| Assembly software used | SPAdes v. 3.9.0 |
| Habitat | Activated sludge |
| Miscellaneous, extraordinary features relevant for the description | - |

**Table S18. Protologue Table for *Candidatus* Propionivibrio dominans**

| | |
|---|---|
| Species name | *Candidatus* Propionivibrio dominans |
| Genus name | *Candidatus* Propionivibrio |
| Specific epithet | dominans |
| Type species of the genus | *Propionivibrio dicarboxylicus* |
| Genus status | Candidatus |
| Species etymology | Description of "*Candidatus* Propionivibrio dominans" sp. nov.: "*Candidatus* Propionivibrio dominans," (do´mi.nans L. masc. adj. dominans, dominant; indicating the high abundance according to 16S rRNA amplioon sequencing). This taxon is represented by the MAG EsbW_MAXAC.044 |
| Species status | sp. nov. |
| Designation of the type MAG | GCA_900099695.1 |
| MAG/SAG accession number | GCA_016709335.1 |
| Genome status | High-quality draft |
| Genome size | 4421984 |
| GC mol % | 58.50 |
| Country of origin | Denmark |
| Region of origin | Esbjerg West |
| Source of sample | Full-scale biological nutrient removal wastewater treatment plant |
| Geographical location | Esbjerg West |
| Latitude | 55.488097 |
| Longitude | 8.430505 |
| Depth | - |
| Altitude | - |
| Temperature of the sample | Mesophilic |
| pH of the sample | - |
| Relationship to oxygen | Facultative anaerobe |
| Energy metabolism | Glycogen-accumulating organism |
| Assembly | 1 sample |
| Sequencing technology | Oxford Nanopore and Illumina Hiseq X |
| Binning software used | MetaBAT2 |
| Assembly software used | CANU v1.8 |
| Habitat | Full-scale biological nutrient removal wastewater treatment plant |
| Miscellaneous, extraordinary features relevant for the description | Rod-shaped cell, with size 0.5-0.6 × 0.9-1.1 µm |

**Table S19. Protologue Table for *Candidatus* Proximibacter danicus**

| | |
|---|---|
| Species name | *Candidatus* Proximibacter danicus |
| Genus name | *Candidatus* Proximibacter |
| Specific epithet | danicus |
| Type species of the genus | *Candidatus* Proximibacter daniensis |
| Genus status | Candidatus |
| Species etymology | Description of "*Candidatus* Proximibacter danicus" gen. nov.: "*Candidatus* Proximibacter danicus," (pro.xi.mi.bac´ter, from L. masc. adj. proximus, near, and N.L masc. n. bacter, bacterium, N.L masc. n. Proximibacter, indicating a rod-shaped bacterium always found attached to filamentous bacteria; da'ni.cus. L. masc. adj. danicus, Danish). This taxon is represented by the MAG Hirt_BATAC.395. |
| Species status | sp. nov. |
| Designation of the type MAG | GCA_016710885.1 |
| MAG/SAG accession number | GCA_016710885.1 |
| Genome status | High-quality draft |
| Genome size | 3246725 |
| GC mol % | 58.60 |
| Country of origin | Denmark |
| Region of origin | Hirtshals |
| Source of sample | Full-scale enriched biological phosphorus removal wastewater treatment plant |
| Geographical location | Hirtshals |
| Latitude | 57.577275 |
| Longitude | 9.992971 |
| Depth | N/A |
| Altitude | N/A |
| Temperature of the sample | Mesophilic |
| pH of the sample | N/A |
| Relationship to oxygen | - |
| Energy metabolism | - |
| Assembly | 1 sample |
| Sequencing technology | Oxford Nanopore and Illumina Hiseq X |
| Binning software used | MetaBAT2 |
| Assembly software used | CANU v1.8 |
| Habitat | Full-scale enriched biological phosphorus removal wastewater treatment plant |
| Miscellaneous, extraordinary features relevant for the description | Rod-shaped cells with size 0.3-05 × 1-2 µm, often attached to filamentous bacteria |
